# Interpretable Atomistic Prediction and Functional Analysis of Conformational Ensembles and Allosteric States in Protein Kinases Using AlphaFold2 Adaptation with Randomized Sequence Scanning and Local Frustration Profiling

**DOI:** 10.1101/2024.02.15.580591

**Authors:** Nishank Raisinghani, Mohammed Alshahrani, Grace Gupta, Hao Tian, Sian Xiao, Peng Tao, Gennady Verkhivker

**Affiliations:** Keck Center for Science and Engineering, Graduate Program in Computational and Data Sciences, Schmid College of Science and Technology, Chapman University, Orange, CA 92866, United States of America; Department of Biomedical and Pharmaceutical Sciences, Chapman University School of Pharmacy, Irvine, CA 92618, United States of America; Department of Pharmacology, Skaggs School of Pharmacy and Pharmaceutical Sciences, University of California San Diego, 9500 Gilman Drive, La Jolla, CA 92093, United States of America; Department of Chemistry, Center for Research Computing, Center for Drug Discovery, Design, and Delivery (CD4), Southern Methodist University, Dallas, Texas, 75275, United States of America

**Author notes:** Correspondence; (G.V).

## Abstract

The groundbreaking achievements of AlphaFold2 (AF2) approaches in protein structure modeling marked a transformative era in structural biology. Despite the success of AF2 tools in predicting single protein structures, these methods showed intrinsic limitations in predicting multiple functional conformations of allosteric proteins and fold-switching systems. The recent NMR-based structural determination of the unbound ABL kinase in the active state and two inactive low-populated functional conformations that are unique for ABL kinase presents an ideal challenge for AF2 approaches. In the current study we employ several implementations of AF2 methods to predict protein conformational ensembles and allosteric states of the ABL kinase including (a) multiple sequence alignments (MSA) subsampling approach; (b) SPEACH_AF approach in which alanine scanning is performed on generated MSAs; and (c) introduced in this study randomized full sequence mutational scanning for manipulation of sequence variations combined with the MSA subsampling. We show that the proposed AF2 adaptation combined with local frustration mapping of conformational states enable accurate prediction of the ABL active and intermediate structures and conformational ensembles, also offering a robust approach for interpretable characterization of the AF2 predictions and limitations in detecting hidden allosteric states. We found that the large high frustration residue clusters are uniquely characteristic of the low-populated, fully inactive ABL form and can define energetically frustrated cracking sites of conformational transitions, presenting difficult targets for AF2 methods. This study uncovered previously unappreciated, fundamental connections between distinct patterns of local frustration in functional kinase states and AF2 successes/limitations in detecting low-populated frustrated conformations, providing a better understanding of benefits and limitations of current AF2-based adaptations in modeling of conformational ensembles.

## Introduction

The remarkable progress of AlphaFold2 (AF2) technology in the field of protein structure modeling have ushered in a transformative era in structural biology, surpassing what was considered feasible just a few years ago.^1,2^ AF2 utilizes evolutionary information by considering Multiple Sequence Alignments (MSAs) derived from related protein sequences as input to capture conserved contacts among evolutionarily linked sequences. The AF2 methodology incorporates the transformer architecture, featuring self-attention mechanisms that allows to discern long-range dependencies and interactions within protein sequences. A pivotal element within the AF2 architecture is the Evoformer module that enables bidirectional information flow throughout the neural network where through self-consistent dynamic exchange of sequence and structure information the network converges to a robust inference.^1,2^ Self-supervised deep learning models, drawing inspiration from natural language processing (NLP) architectures, and particularly approaches incorporating attention-based^3^ and transformer mechanisms^4^ have proven to be powerful tools for training on extensive sets of protein sequences. The attention mechanism, originally introduced in the context of NLP, allows the model to focus on different parts of the input sequence, accurately capturing the contextual relationships.^3^ Transformer mechanisms, which have excelled in language tasks, contribute to the model’s ability to efficiently handle sequential data.^4^ These self-supervised models have demonstrated their efficiency by learning from large ensembles of protein sequences. Notably, they exhibit the capacity to predict protein structures from individual sequences, eliminating the dependence on MSA information.^6,7^ Among latest breakthrough in AI-based protein structure predictions is a high accuracy end-to-end family of transformer protein language model ESMFold for atomic level structure prediction directly from the individual sequence of a protein.^6^ OmegaFold represents another related approach and utilizes a hybrid model that combines a protein language model with a geometry-guided transformer model that is trained on protein structures, enabling robust predictions from individual sequences.^7^ Unlike AF2, OmegaFold contains a protein language model that processes the full sequence to generate a latent space which serves as input to a GeoFormer network incorporating knowledge of protein geometry and topology to create a final prediction. OmegaFold outperforms RoseTTAFold approach^8^ and achieves a comparable level of prediction accuracy to AF2 without relying on MSA information. RoseTTAFold2 approach extends the original three-track architecture of RoseTTAFold^8^ over the full network by integrating AF2 mechanism of updating pair features using a more computationally efficient structure-biased attention, producing comparable accuracy on monomers, and AF2-multimer on complexes while allowing for efficient computational scaling on large proteins and complexes.^9^ Recent study from Baker laboratory employed variational autoencoders (VAE) trained on the crystal structures and simulation snapshots to convert protein structural data into a continuous, low-dimensional representation, followed by guided sampling in the latent space and rapid generation of accurate conformational ensembles with RoseTTAFold.^10^ A new method for accurate protein design RoseTTAFold Diffusion integrates structure prediction networks and diffusion generative model enabling design of complex functional proteins from simple molecular specifications.^11^ Different from the AF2, the trained language models can learn directly evolutionary patterns of sequences linked to structure, eliminating the need for MSAs and templates which improves the speed of protein structure prediction while maintaining high resolution accuracy comparable to AF2 models. The success of these approaches highlights the evolving landscape of techniques in the field, displaying how advancements in language models and direct learning from sequences can lead to substantial improvements in the speed and accuracy of protein structure prediction tools.

Despite the remarkable success of AF2-based methods and self-supervised protein language models that exceled at predicting static protein structures, there are notable shortcomings related to their applicability and generality in accurately characterizing conformational dynamics, functional protein ensembles, conformational changes, and allosteric states.^12^ Conformational changes and allosteric states involve subtle shifts in protein structure and function and are crucial for understanding how proteins interact with other molecules and modulate cellular processes. Several recent studies indicated that protein structure prediction capabilities of the AF2 methods are not trivially expandable for prediction of conformational ensembles and accurate mapping of allosteric landscapes.^13–17^ Efforts to optimize the AF2 methodology for predicting alternative conformational states of proteins have primarily centered around manipulating the MSA information which is motivated by the recognition that MSAs should encode for coevolutionary signals of the most thermodynamically probable protein state and also for other functional conformational states of a protein.^13,14^ These approaches assume that by efficiently deconvolving and interpreting these signals, AF2 method can be modified and adapted to enable prediction single dominant conformation and conformational ensembles. In one of these approaches, MSAs are randomly subsampled to reduce depth, resulting in shallower MSAs which aims to enhance the diversity and produce a broader range of the AF2 output models to capture experimentally validated alternative conformational states of proteins.^13^ The alternative AF2-based approach known as SPEACH_AF (Sampling Protein Ensembles and Conformational Heterogeneity with AlphaFold2) involves the manipulation of the MSA through in silico mutagenesis where replacing specific residues within the MSA can induce changes in the distance matrices, ultimately leading to the prediction of alternate protein conformations.^14^ In this approach, it is assumed that alanine mutations in the MSA could broaden an attention network mechanism within AF2 and ascertain distinct patterns of coevolved residues associated with alternative conformations.^14^

The limitations of AF2 methods in predicting multiple protein conformations may be associated with the intrinsic training bias for the experimentally determined, thermodynamically stable structures and MSAs containing the evolutionary information used to infer the ground protein states. In particular, the AF2 methodology can be systematically biased to predict only one conformation of fold-switching proteins as 94% of the AF2 predictions captured the experimentally determined ground conformation but not the alternative structure.^16^ The latest study from the same laboratory examined more systematically the AF2 adaptations for sampling of alternative states for predictions of experimental conformational states by generating >280,000 models of 93 fold-switching proteins with experimentally determined conformations.^17^ Combining all models, the AF2 predicted structures displayed only modest success and were often unable to detect the experimentally characterized conformations for most fold switching proteins. Moreover, by generating >159,000 models on seven fold-switching proteins outside of the training set, AF2 yielded accurate fold switching predictions only in one of seven targets with moderate confidence.^17^

Some of the recent attempts for expanding AF2 capabilities towards prediction of conformational ensembles involved combination of shallow MSA and state-annotated templates incorporating functional or structural properties of GPCRs and protein kinases.^18^ Another AF2 adaption termed AF-Cluster used a simple MSA subsampling method for subsequent clustering of evolutionarily related or functionally similar sequences, enabling predictions of alternative protein states and showing promise in identifying previously unknown fold-switched state that was later validated by the NMR analysis.^19^ Recent application studies indicated that AF-cluster predicts fold switching from single sequences by associating structures in its training set with homologous sequences, suggesting that without coevolutionary information, AF-cluster may be constrained to sample alternative conformations available during training and therefore may have limited applicability for novel fold-switching proteins that are absent in the training set.^20^ Cfold is a recently proposed structure prediction network approach that is similar to AF2 but is trained on a conformational split of the Protein Data Bank (PDB) for predicting alternative conformations of protein structures by using dropout and MSA clustering tools.^21^ Several user-friendly powerful architectural pipelines that facilitate AF2 implementations were developed including ColabFod^22^ and OpenFold.^23^ OpenFold is a memory-efficient and trainable open-source implementation of AF2 with OpenProteinSet, a database of five million deep and diverse MSAs, allows for faster prediction of large proteins and multi-protein complexes. OpenFold^23^ and Uni-Fold^24^ are retrainable implementations of AF2 that can integrate the experimental data derived from crosslinking experiments to guide monomeric and multimeric predictions. Another study introduced AF unmasked adaptation that can leverage information from templates containing quaternary structures without the need for retraining and can generate robust models of large complexes in one shot allowing for robust integrative structure modeling and predictions of protein dynamics.^25^

The analysis of various AF2 pipelines reveals a significant focus of emerging adaptations on broadening the range of accessible protein conformations to enhance the ability to predict specific conformational states by adjusting the balance between genetic information obtained from MSAs and structural information derived from templates.^26^ Expanding existing tools for prediction of allosteric regulatory mechanisms would require a considerable step forward towards accurate mapping of the conformational ensembles and prediction of functional allosteric conformations.^27^ Recent analysis of AF2 predictions and direct comparison with the experimental crystallographic maps showed that in many cases even high-confidence predictions could substantially differ from the experimental maps through distortions and domain orientations, and on a local scale in the backbone and side-chain conformation, indicating that experimental structural determination remains the key step in validating the AF2-based hypotheses and computational predictions.^28^

Protein kinases that are regulatory machines characterized by the multiple functional states of the catalytic domain^29–34^ present a class of regulatory switches that operate through dynamic equilibrium between the active and inactive states. Despite significant structural wealth of the protein kinases and their complexes with ATP-bound inhibitors and allosteric modulators^35,36^, the atomistic characterizations of the functionally relevant transient states has been lacking due to large conformational transformations and short-lived kinase intermediates involved in kinetics of the allosteric shifts. The recent NMR-based structural determination of the unbound ABL kinase in the active state and two inactive functional conformations that are unique for ABL kinase presents an ideal challenge for AF2 approaches (Figure 1).^37,38^ The thermodynamically dominant active conformation of the ligand-free ABL kinase and two short-lived inactive conformations I_1_ and I_2_ that that occur only 5% of the time are intrinsically present on the conformational landscape of the unbound kinase domain and are quite different from each other in the critical functional regions (Figure 1).^38^ Conformational transitions between the inactive and active kinase states are orchestrated by three conserved structural motifs in the catalytic domain: the αC-helix, the DFG-Asp motif (DFG-Asp in, active; DFG-Asp out, inactive), and the activation loop (A-loop open, active; A-loop closed, inactive) providing structural fingerprints that differentiate the active and inactive forms (Figure 1).

**Figure 1.**
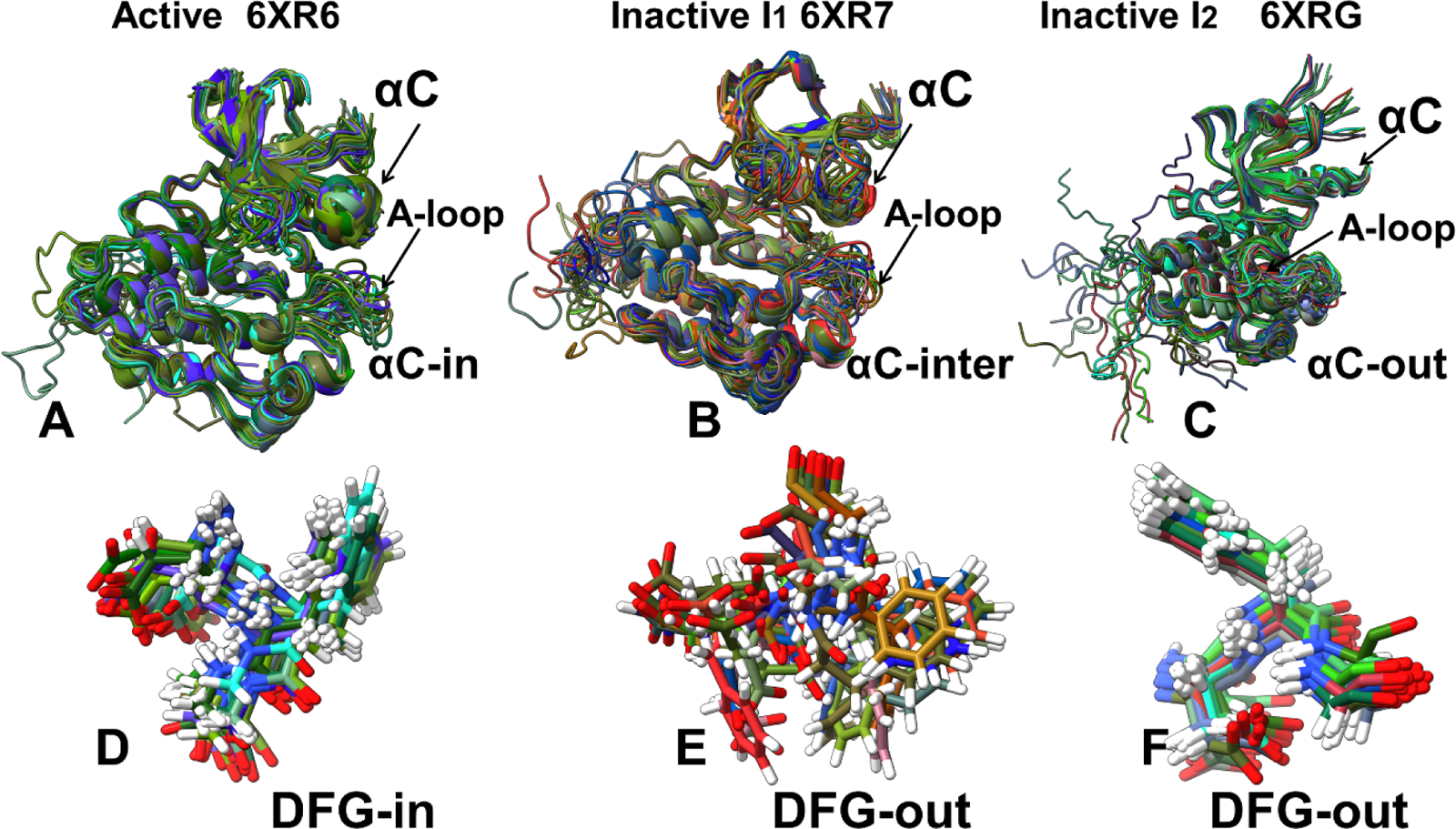
The NMR solution structural ensembles of the thermodynamically stable fully active ground state of the ABL kinase domain (pdb id 6XR6) (A), the inactive state I_1_ (pdb id 6XR7) (B) and the closed inactive state I_2_ (pdb id 6XRG) (C). The NMR ensemble of DFG conformations in the active ABL state (D), the inactive state I_1_ (E) and the closed inactive state I_2_ (F). The ABL conformations are shown in ribbons. The structures point to similarities and differences in the key functional regions of the kinase domain exemplified by the αC helix, the A-loop, and the P-loop. In particular, A-loop in the inactive state I_2_ (C,F) adopts a completely different closed conformation.

Multidisciplinary studies that exploited synergies between structural, biophysical approaches and computational methods^40–42^ have been fruitful in uncovering the invisible dynamic aspects of protein kinases. Our recent study employed atomistic molecular dynamics (MD) simulations with dimensionality reduction methods and Markov state models (MSM) to characterize the dynamics and kinetics of structural changes between the ABL conformational states.^42^ While MD simulations and MSM provided insights into conformational dynamics of the NMR-determined ABL states, these approaches completely relied on the initial experimental structures. A far more challenging problem is to computationally predict the ABL structures including the low-populated inactive conformations and determine conformational ensembles of these states without a priori experimental knowledge of these states. Several recent investigations explored the potential of AF2 methodologies for predicting conformational states in protein kinases. AF2-based modeling of 437 human protein kinases in the active form using shallow MSAs of orthologs and close homologs of the query protein showed the robustness of AF2 methods as selected models for each kinase based on the prediction confidence scores of the activation loop residues conformed closely to the substrate-bound experimental structures. ^43^ The ability of AF2 methods to predict kinase structures in different conformations at various MSA depths was examined, demonstrating that using lower MSA depths allows for more efficient exploration of alternative kinase conformations, including identification of previously unseen conformations for 398 kinases.^44^ Another AF2 adaptation explored the conformational landscape of the ABL kinase domain by systematically manipulating the MSA subsampling parameters and predicted the relative populations of different ABL conformations.^45^ While this study suggested that AF2 with shallow MSAs may be useful for predicting conformational ensembles in protein kinases, accurate and reproducible prediction of functional allosteric conformations in the ABL kinase, particularly low-populated inactive states, remains challenging and is highly sensitive to the architectural details of the AF2 pipeline.

In the current study, we employ and systematically compare several recent AF2 adaptations including (a) MSA subsampling with shallow MSA depth; (b) SPEACH_AF approach in which alanine scanning is performed on generated MSAs by using different random alanine mutation positions in the MSAs; and (c) random alanine scanning algorithm that iterates through each amino acid in the native sequence and randomly substitutes 5-15% of the residues to simulate random alanine substitution mutations. The results demonstrate that the proposed AF2 adaptation enables accurate and robust prediction of structural ensembles and conformational heterogeneity of the active and the intermediate inactive ABL state that represents one of the two common inactive forms found in the crystal structures of protein kinases.^46,47^ We show that the proposed AF2 adaptation combined with local frustration mapping of conformational states enable accurate prediction of the ABL active and intermediate structures and conformational ensembles, also offering a robust approach for interpretable characterization of the AF2 predictions and limitations in detecting hidden allosteric states. We show that the dominant minimal frustration pattern in the active ABL state and the inactive intermediate I_1_ state provide a broad and funnel-liked landscapes around these states allowing AF2 predictions to accurately capture structural ensembles of these functional ABL conformations. In contrast, the emergence of interconnected high frustration residue clusters in the inactive I_2_ state that define the initiation “cracking” sites of allosteric changes and present difficult targets for robust AF2 predictions. This study uncovered previously unappreciated, fundamental connections between distinct patterns of local frustration in functional kinase states and AF2 successes/limitations in detecting low-populated frustrated conformations, providing understanding and rationale for limitations of current AF2-based adaptations in modeling of conformational ensembles.

## Materials and Methods

### Protein Structure Modeling Using AF2 with MSA Shallow Subsampling Adaptation

Structural prediction of the ABL kinase states were carried out using AF2 framework^1,2^ within the ColabFold implementation^22^ using a range of MSA depths and MSA subsampling.^13^ We used *max_msa* field to set two AF2 parameters in the following format: *max_seqs:extra_seqs*. These parameters determine the number of sequences subsampled from the MSA (*max_seqs* sets the number of sequences passed to the row/column attention track and *extra_seqs* the number of sequences additionally processed by the main evoformer stack). The lower values encourage more diverse predictions but increase the number of misfolded models. The default MSAs are subsampled randomly to obtain shallow MSAs containing as few as five sequences (Figure 2). We set the *max_msa* parameter to 16:32. This parameter is in the format of *max_seqs:extra_seqs* which decides the number of sequences subsampled from the MSA. *Max_seq* determines the number of sequences passed to the row/column attention matrix at the front end of the AF2 architecture, and *extra_seqs* sets the number of extra sequences processed by the Evoformer stack after the attention mechanism. We additionally manipulated the *num_recycles* parameters to produce more diverse outputs. We use *num_recycles*: 12. AF2 makes predictions using 5 models pretrained with different parameters, and consequently with different weights. To generate more data, we set the number of recycles to 12, which produces 14 structures for each model starting from recycle 0 to recycle 12 and generating a final refined structure. Recycling is an iterative refinement process, with each recycled structure getting more precise. Each of the AF2 models generates 14 structures, amounting to 70 structures in total (Figure 2). In addition, we also predicted ABL structure using AF2 with the default and ‘auto’ parameters serving as a baseline structure for prediction and variability analysis.

**Figure 2.**
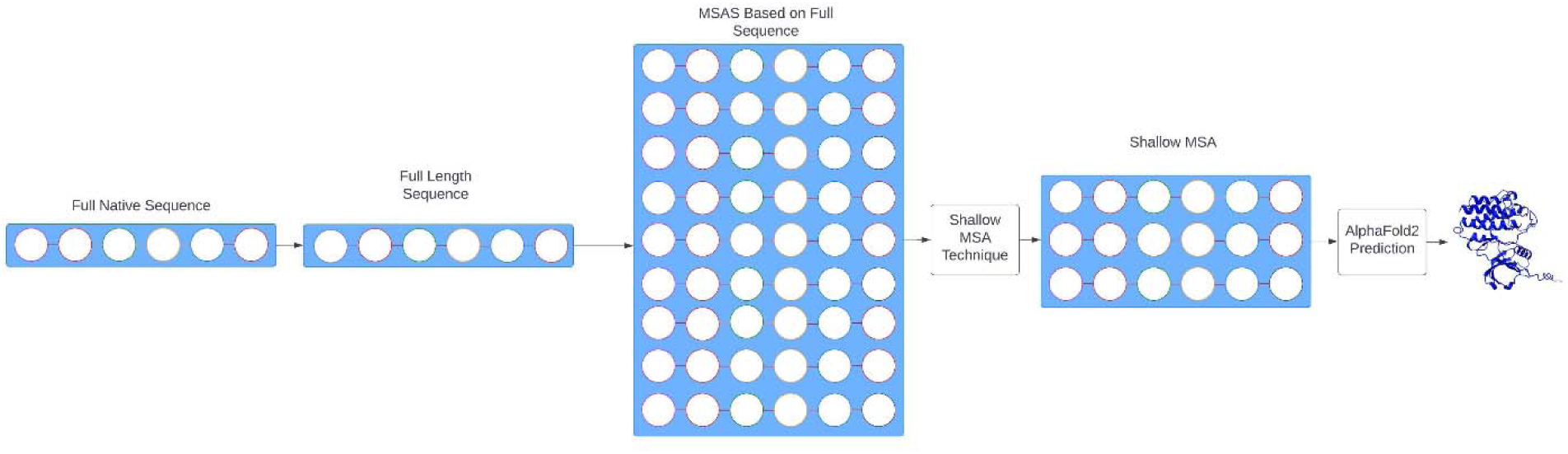
A schematic representation of the AF2 protein structure prediction pipeline using MSA shallow subsampling.

The MSAs were generated using the MMSeqs2 library using the ABL1 sequence from residues 240 to 440 as input. We then set the *num_seed* parameter to 1. This parameter quantifies the number of random seeds to iterate through, ranging from *random_seed to random_seed+num_seed*. Increasing the *num_seeds* samples predictions from the uncertainty of the model. We also enabled the use of dropout parameter, meaning that dropout layers in the model would be active during the time of predictions, which further increases variability within predictions. As a summary of the process, we input a protein sequence of the ABL1 kinase and predicted 70 unique structures through the shallow MSA technique.

### Protein Structure Modeling Using AF2 with SPEACH AF Adaptation

In SPEECH_AF approach^14^, MSAs were first generated from the original full-length protein sequence using MMSeqs2. The method then applied random alanine masking to the generated MSA. This introduced alanine mutations at the same positions in all homologous sequences within the MSA. Differing from our other approach of alanine scanning the original sequence prior to MSA construction, this method generates the MSA first and then performs alanine scanning. We generated multiple distinct MSAs using the SPEACH AF notebook, by using different random alanine mutation positions in the MSAs as well as the amount of mutations in the MSAs. We chose two MSAs out of these, with the first MSA also having a lower frequency of mutation than the second one (Figure 3).

We start with native sequence S = {r_0_, r_1_, r_2_,…, r_n_}, which we use to create an MSA.

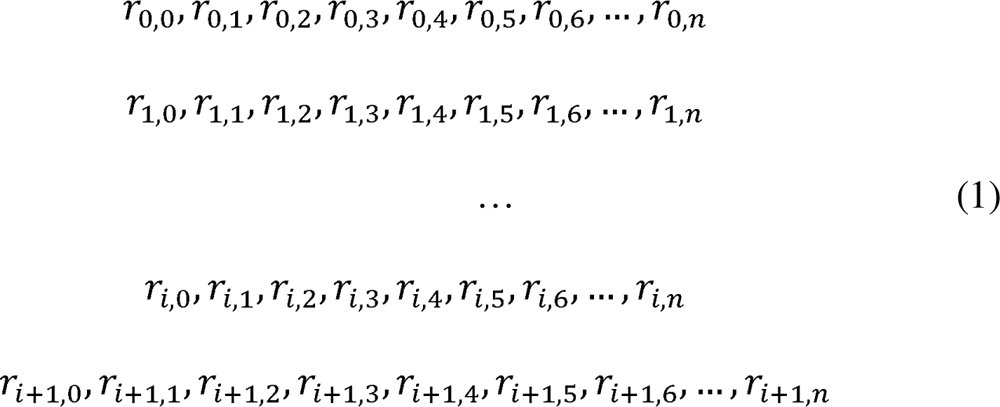

This MSA is then processed by SPEACH AF approach where it undergoes alanine scanning of all sequences in the MSA, from which we take two distinct MSA sequences.

MSA 1:

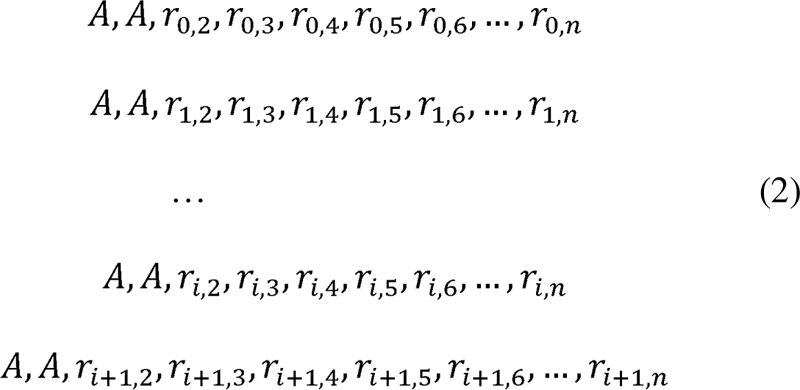

MSA 2:

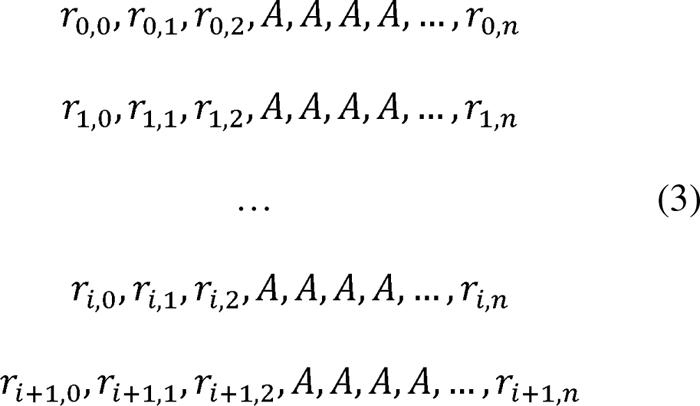

**Figure 3.**
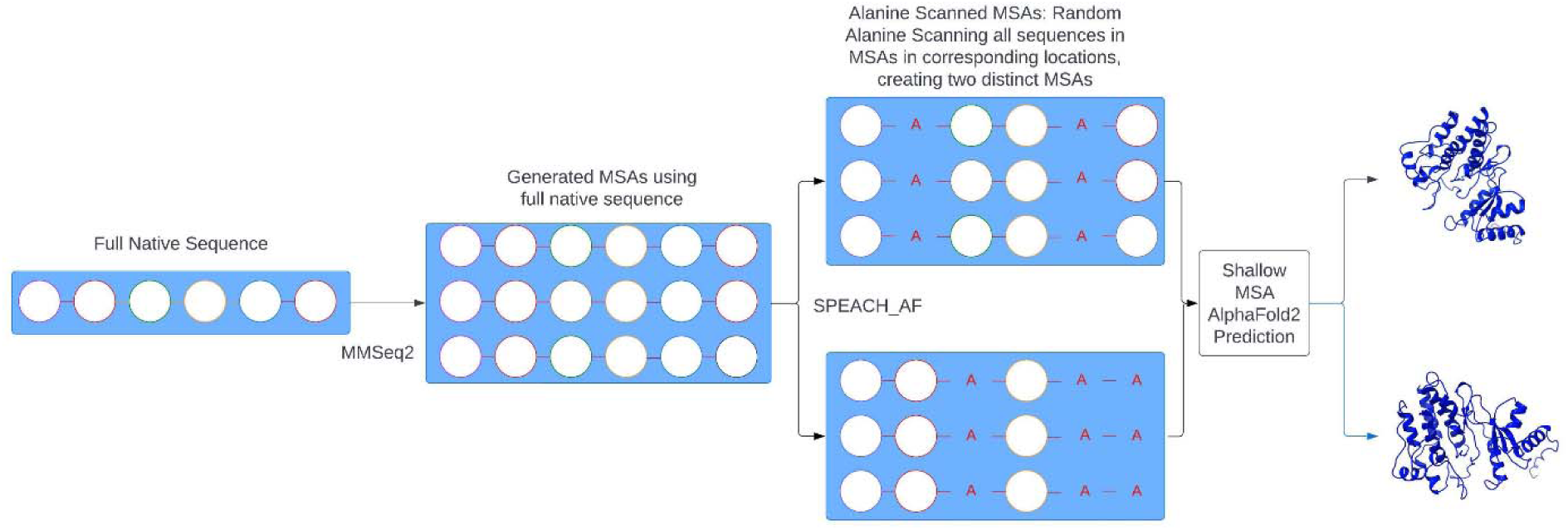
A schematic representation of the AF2 protein structure prediction pipeline using SPEACH_AF adaptation.

This technique attempts to limit the bias toward the dominant native sequence form that may occur when constructing the MSA after alanine scanning of the full sequence. The generated MSAs were custom MSA inputs for AF2. The shallow MSA methodology was then employed to make predictions and further increase structural variability. The predicted structures were generated from 6 recycles per model.

### Protein Structure Modeling Using Randomized Alanine Sequence Scanning and MSA Shallow Subsampling for Generation of Conformational Ensembles

The initial input for the full sequence randomized alanine scanning (Figure 4) is the original full native sequence. This technique utilizes an algorithm that iterates through each amino acid in the native sequence and randomly substitutes 5-15% of the residues with alanine, to simulate random alanine substitution mutations. The algorithm substitutes residue a_i_ with alanine at each position i with a probability p_i_ randomly generated between 0.05 and 0.15 for each sequence position. We ran this algorithm nine times on the full native sequence, resulting in nine distinct sequences, each with different frequency and position of alanine mutations (Figure 4). Multiple sequence alignments (MSAs) were then constructed for each of these mutated sequences using the alanine-scanned full-length sequences as input for the MMSeqs2 program.

We start with native sequence S = {r_0_, r_1_, r_2_,…, r_n_}, which we then apply the alanine mutation algorithm to resulting in nine new sequences.

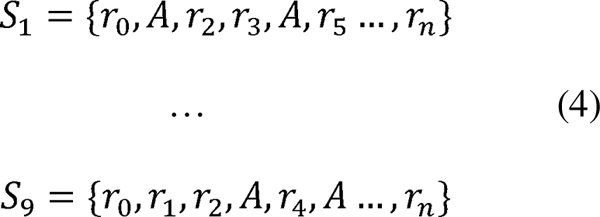

We then use these full sequences to generate our MSAs.

MSA 1:

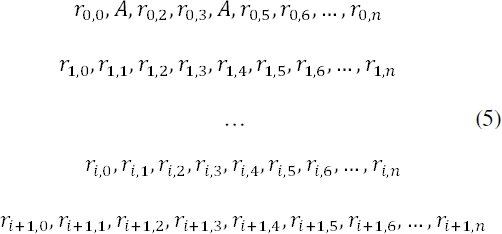

MSA 9:

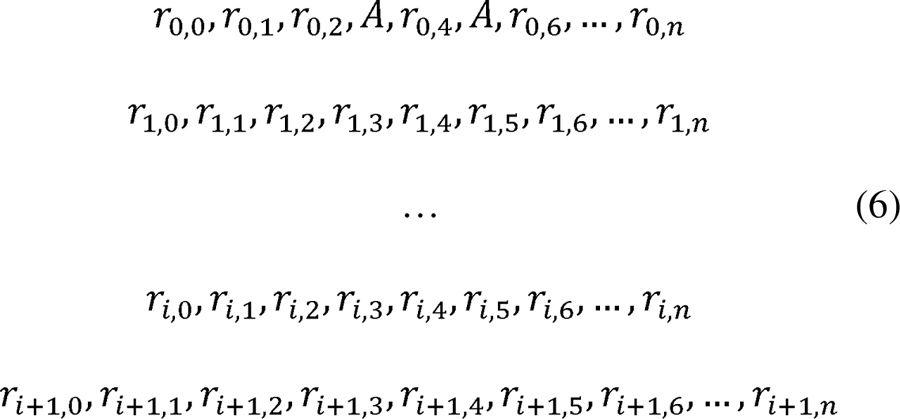

**Figure 4.**
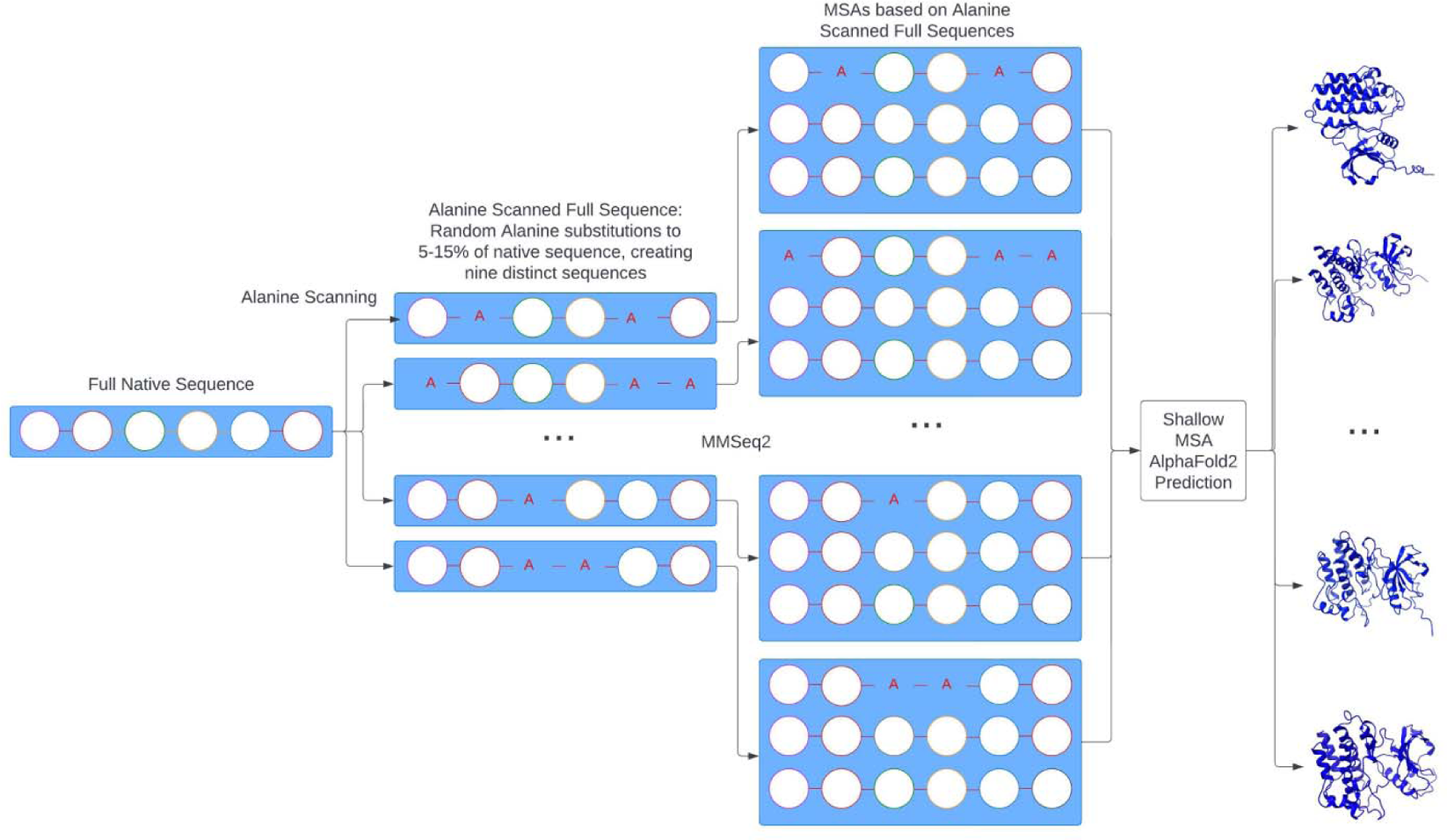
A schematic representation of the AF2 protein structure prediction pipeline using full sequence randomized alanine scanning and MSA shallow subsampling.

This method aims to perturb the MSAs without actually mutating the homologous sequences within them, which is the major difference from the SPEACH_AF approach. The AF2 shallow MSA methodology is subsequently employed on these MSAs to predict protein structures as described previously. A total of 70 predicted structures were generated from 12 recycles per model.

### Integration and Comparative Analysis of AF2 Adaptations with Shallo MSA Depth and Alanine Mutagenesis Scanning of MSAs and Sequences

To develop accurate and robust atomistic models of the ABL structure and dynamics, we performed comparative structural prediction using a hierarchy of several AF2-based adaptations : AF2 default settings, AF2 methodology with shallow MSA depth^13^ ; SPEACH_AF in which the MSAs are manipulated via *in silico* mutagenesis by replacing specific residues within the MSAs^14^ ; and random alanine scanning that iterates through each amino acid in the native sequence and randomly substitutes the residues to simulate random alanine substitution mutations (Supplementary Information, Figure S1). By systematically applying and comparing these three AF2 pipelines along with the manipulation of the AF2 parameters, we predict the structures and populations of the ABL conformational ensembles. In its standard implementation, AF2 takes as input a target sequence and a corresponding multiple sequence alignment. An arbitrary number of sequences (defined by the *max_seq* parameter) are randomly selected from the master MSA (the target sequence is always selected), and the remaining sequences are clustered around each of the selected sequences using a Hamming distance. Through analysis of the AF2-based adaptations which shared as their core subsampling of the MSAs, we identified how different methods affect the prediction quality and reliability of AF2 tools. By varying AF2 parameters, we examined whether a systematic reduction of the MSA depth can alter conformational diversity of the ABL kinase models. We optimized the accuracy achieved as a function of the following parameters: *max_seq, extra_seq*, number of seeds, and number of recycles. We generated five models for each kinase, which were then classified into the same six conformations.

### Statistical and Structural Assessment of AF2-Generated Models

AF2 models were ranked by Local Distance Difference Test (pLDDT) scores (a per-residue estimate of the prediction confidence on a scale from 0 to 100), quantified by the fraction of predicted Cα distances that lie within their expected intervals. The values correspond to the model’s predicted scores based on the lDDT-Cα metric which is a local superposition-free metric that assesses the atomic displacements of the residues in the predicted model.^1,2^

Models ranked in the top five were compared to the experimental structure using structural alignment tool TM-align, an algorithm for sequence-independent protein structure comparison, to assess and compare the accuracy of protein structure predictions.^48^ TM-align involves optimizing the alignment of residues, refining the alignment through dynamic programming iterations, superposing the structures based on the alignment, and finally, calculating the TM-score as a quantitative measure of the overall accuracy of the predicted models. This approach provides a systematic and quantitative way to evaluate and rank the structural accuracy of predicted models in comparison to experimentally determined reference structures. TM-align calculates the TM-score as the measure of overall accuracy of the prediction. An optimal superposition of the two structures is then built and TM-score is reported as the measure of overall accuracy of prediction for the models. TM-score ranges from 0 to 1, where a value of 1 indicates a perfect match between the predicted model and the reference structure. When TM-score > 0.5 implies that the structures share the same fold. TM-score > 0.5 is often used as a threshold to determine if the predicted model has a fold similar to the reference structure. If the TM-score is above this threshold, it suggests that the predicted structure and the reference structure have a significant structural resemblance. Several other structural alignment metrics were used including the global distance test total score GDT_TS of similarity between protein structures and implemented in the Local-Global Alignment (LGA) program^49^ and the root mean square deviation (RMSD) superposition of backbone atoms (C, Cα, O, and N) calculated using ProFit (http://www.bioinf.org.uk/software/profit/). Distributions of TM-Scores were calculated in terms of the averages of the highest TM-Scores for AF2 models compared to PDB structures of the same kinases for each MSA depth and structural conformation.

### Local Frustration Analysis

We modeled local frustration by computing a residue-based local frustration index for the structural ensembles obtained from AF2 predictions via a web server (http://www.frustratometer.tk).^50–22^ We quantified conformational and mutational frustration of protein residues. This analysis is based on the profiling of the protein kinase residues by local frustratometer^50–52^ which computes the local energetic frustration index using the contribution of a residue to the energy in a given conformation as compared to the energies that would be found by mutating residues in the same native location or by creating by changing local conformational environment for the interacting pair.^50–52^ The local frustration index for the contact between the amino acids i,j was defined as a Z-score of the energy of the native pair compared to the N decoys. A residue-based frustration index measures the energetic stability of a particular native contact as compared to a set of all contacts sampled by generating a large number of distributed decoys and recomputing the energy change. For mutational frustration, the decoy set was made randomizing the identities of the interacting amino acids i, j, keeping all other interaction parameters at their native value. This scheme effectively evaluates every mutation of the amino acid pair that forms a particular contact in a fixed structure. For configurational frustration, the decoy set involves randomizing not only the residue identities but also the distance between the interacting amino acids i, j. The frustration index was calculated by mutating the identities and the distance between the interacting amino acids. The index value that corresponded to a positive Z-score value would indicate that the majority of other amino acid pairs in that position were unfavorable. In this approach, a significant stabilization for an individual native pair normalized by the energy fluctuations is considered as indication of minimally frustrated interaction, whereas a destabilizing effect of the native pair deems the corresponding interaction as highly frustrated.

To quantify the role of molecular frustration in the allosteric control of ABL kinase and evaluate local frustration patterns the active and inactive ABL states, we employed FrustratometeR appraratus.^51,52^ We evaluate configurational and mutational molecular frustration by computing local densities of minimal, neutral and high frustration contacts within 5 Å sphere from a given protein residue. The local frustration analysis is conducted on the NMR-determined ABL structures and the AF2-predicted ABL conformations from the determined structural ensembles. The distributions of neutral, high and minimal frustration are determined and compared with a more general frustration analysis that revealed the majority of local interactions as neutral (∼50-60%) or minimally frustrated ( 30%) with only 10% of the total contacts considered as highly frustrated.^50^ The local frustration analysis is interpreted in the context of frustration surveys of the diverse protein folds showed that minimally frustrated residues may be enriched in the protein core, while the exposed flexible binding interfaces are characterized by the highly frustrated patches that can be alleviated upon binding.^55^ The local frustration patterns allow for detection of conserved frustration hotspots associated with kinase regulatory function to orchestrate conformational transitions between the active and inactive states. By combining AF2 predictions of kinase states with local frustration mapping of conformational states we propose a model for interpretable characterization and assessment of AF2 predictions.

## Results and Discussion

### Structural Analysis of the NMR Conformational Ensembles of the ABL Kinase

First, we analyzed in more detail the NMR-generated ensemble of the ligand-free ABL kinase domain^38^ in its active conformational state (pdb id 6XR6) and two inactive conformational states I_1_ and I_2_ (pdb id 6XR7, 6XRG) (Figure 1). Structural analysis showed that the NMR ensemble of the active conformation (Figure 1A) closely conforms to this thermodynamically dominant state with some appreciable degree of conformational plasticity that is visible in the A-loop and the αC helix. Nonetheless, all conformations from the active ensemble are characterized by the “αC-in” position and stable DFG-in orientation (Figure 1A,D). In the active “αC-in” state a conserved αC helix residue E305 forms an ion pair with K290 in the β3 strand that coordinates the α and β phosphates of the ATP. Interestingly, the NMR ensembles of the inactive ABL states are very different structurally and dynamically (Figure 1B,C). In the inactive I_1_ state (pdb id 6XR7) the αC helix moves from its active “αC-in” position to the intermediate αC-in/out position in the I_1_ state (Figure 1B,E). The structural alignment of the DFG conformations in the I_1_ ensemble highlighted a considerable variability in the regulatory DFG motif that adopts distinct “out” conformations exemplified by heterogeneity of the phenylalanine departures from the active “in” position (Figure 1E). In the I_1_ ensemble the DFG motif is flipped 180°, with respect to the active conformation, but the A-loop remains in an open and highly heterogeneous conformation similar to the active conformation. In this form, ABL adopts the DFG-out, αC-helix-in conformation and thus it is catalytically inactive (Figure 1B,E) . The analysis of the NMR ensembles showed a contrast between heterogeneous I_1_ ensemble (Figure 1B,E) and a more restricted inactive I_2_ ensemble which is different structurally and dynamically (Figure 1C,F). The regulatory DFG motif adopts a distinct “out” conformation in the I_2_ state (Figure 1F) where only minor displacements were observed in the NMR ensemble of this inactive state. In this inactive form the A-loop undergoes a large conformational arrangement to adopt fully closed conformation which is accompanied by change in the αC-out position (Figure 1C,F). Similar to the I_1_ state, I_2_ adopts the DFG-out conformation. F401 of the DFG in the I_2_ state flips into the catalytic pocket and translates by more than 10 Å to occupy a hydrophobic pocket that is lined up by L267, V275, A288, F336, and L403 residues (Figure 1C,F). Interestingly, the original structural studies remarked that while Abl I_2_ state adopts a fully inactive conformation, the inactive intermediate I_1_ state may lie in the pathway between the active and the I_2_ state.^38^ Moreover, it was suggested that the I_1_ state may have evolved during the evolutionary process as a byproduct that established the active and the fully inactive I_2_ state, as the two primary functional states.^38^

### AF2 Atomistic Modeling of the Conformational Ensembles of the ABL Kinase : Shallow MSA Depth Increases the Structural Diversity of AF2-Generated Kinase Models

The important objective of this study was to explore the potential and limits of various AF2 adaptations while keeping the overall AF2 implementation architecture and MSA generation procedure as was initially prescribed. First, we analyzed the AF2 results using shallow MSA depth settings, which showed accurate predictions of the ABL active state and excellent structural alignment with the crystallographic active conformation with high confidence pLDDT values (Figure 5). Moreover, the top AF2 predicted models selected by pLDDT values displayed increased structural similarity to the experimental ABL1 active kinase structure. The analysis of the pLDDT profiles for the predicted top models showed high confidence values for the most of the ABL kinase domain regions (pLDDT ∼80-100), while the highly mobile A-loop (residues 395-421) displayed variability of pLDDT values (pLDDT ∼65-85) (Figure 5A).

**Figure 5.**
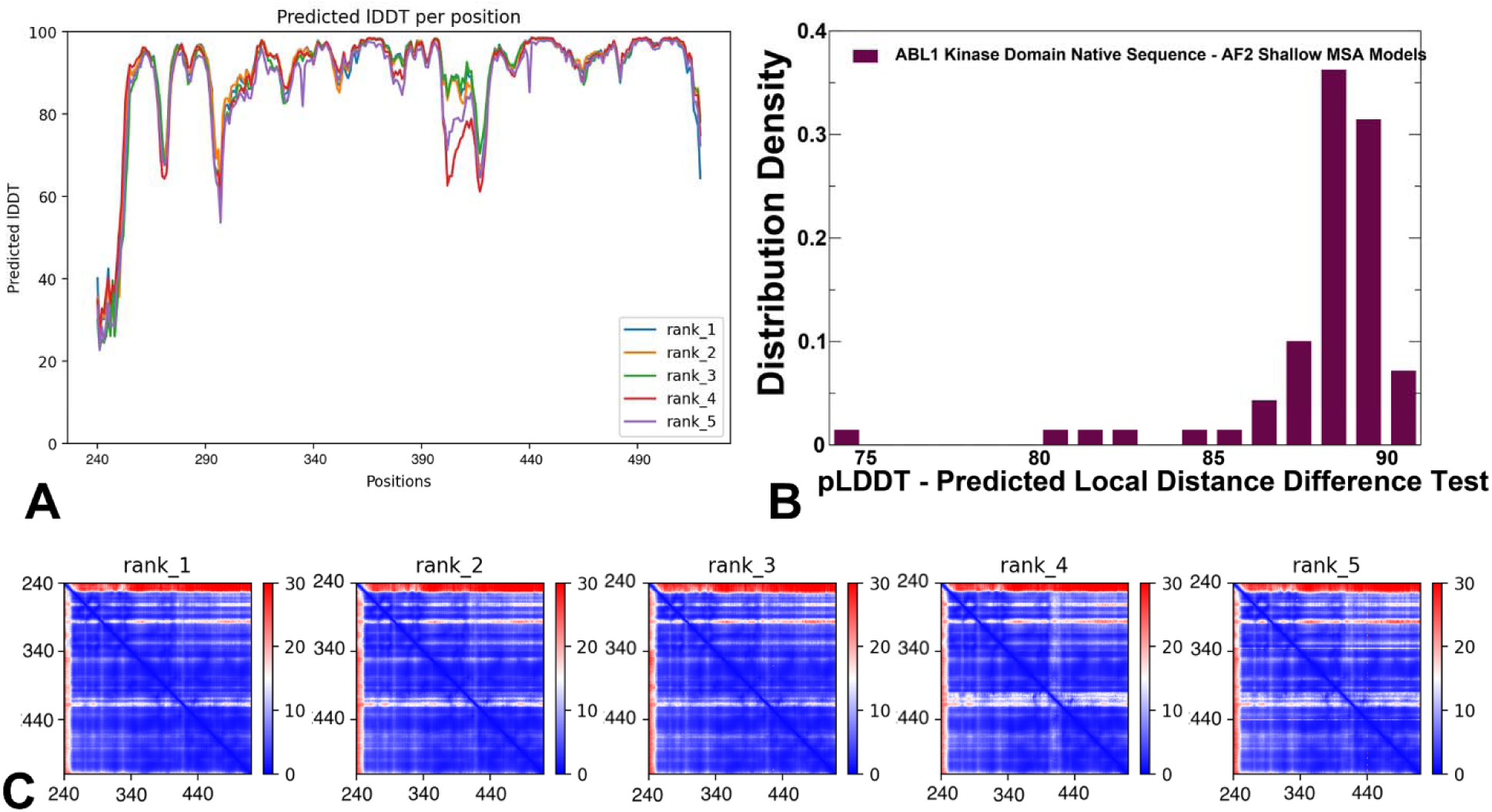
The statistical analysis of the shallow MSA depth models for the ABL kinase structures. The residue-based pLDDT profile for the top five ranked models is obtained from AF2 predictions of ABL conformations (A). The distributions of structural model assessment for the ABL conformational ensemble are obtained from AF2-MSA shallow subsampling predictions. (B) The density distribution of the AF2-derived pLDDT structural model estimate of the prediction confidence on a scale from 0 to 100. The predicted alignment error (PAE) heatmaps for the top five models (C). The heat maps are provided for each top ranked model and show the PAE between each residue in the model. The color scale contains three colors to highlight the contrast between the high confidence regions and the low confidence regions.

In general, protein regions with pLDDT values ∼ 50-70 have lower confidence, while pLDDT < 50 may indicate some level of disorder. Indeed, some of the low confidence pLDDT values corresponded to disordered N-terminal residues that were revealed in the NMR ensembles. The density distribution of the pLDDT values obtained for the AF2 shallow MSA structural ensemble showed a pronounced peak at pLDDT ∼90 and several minor peaks for pLDDT∼80-85 and pLDDT ∼75 (Figure 5B). The heat maps of the PAE for the best AF2 five models (Figure 5C) highlighted differences between the high low confidence regions, pointing to the increased heterogeneity of the A-loop (residues 395-421) in the bottom three models.

We followed up with the evaluation of structural similarity metrics TM scores and RMSD values computed for the complete predicted ABL structure and only for the A-loop residues in the AF2-predicted conformations (Figure 6). Structural similarity of the predicted conformations was evaluated with respect to the active and inactive experimental ABL structures. The TM scores showed the most dominant peak of TM values ∼0.95 with respect to the active ABL conformation, while only a small fraction of the AF2 ensemble was similar to the I_1_ state and the predicted conformations were vastly different from the inactive state I_2_ (Figure 6A). The RMSD distribution for the ABL residues similarly highlighted the peak at RMSD∼0.2-0.4 Å from the active state and another strong peak at RMSD∼1.0 Å from the highly mobile inactive state I_1_ (Figure 6B). By specifically analyzing density distributions of the RMSD for the A-loop residues with respect to the experimental ABL states, we found three peaks at RMSD = 1.0 Å, 2.0 Å and 3.0 Å from the active ABL structure (Figure 6C), thus confirming the ability of the AF2 method not only accurately predict the dominant ABL state but also capture the intrinsic heterogeneity of the open A-loop in the active form. The predicted A-loop conformations also displayed a shallow peak at RMSD∼3.0 Å from the inactive I_1_ state which represented a fraction of the predicted AF2 ensemble conformations that were similar to the I_1_ state, while most of the AF2-predicted structural ensemble conformations were structurally different from the inactive I_2_ state (Figure 6C).

**Figure 6.**
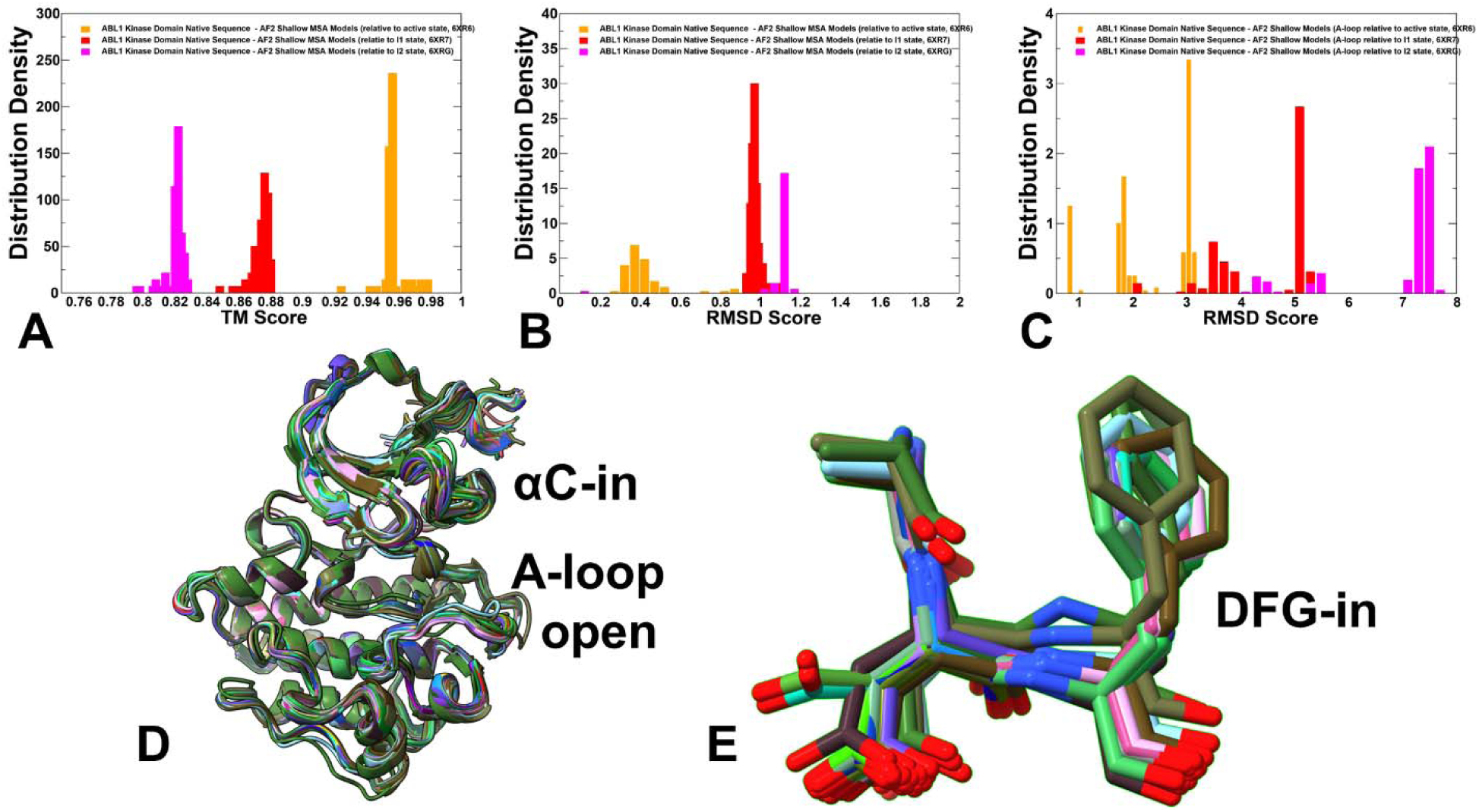
The distribution of structural similarity metrics TM scores and RMSD values for the predicted ABL conformations and for the predicted A-loop conformations computed with respect to the experimental active and inactive ABL states. (A) The distribution density of TM scores for the AF2-predicted ABL conformations using shallow MSA depth with respect to the experimental active ABL state (orange filled bars) and the inactive states I_1_ (red filled bars) and I_2_ (blue filled bars). (B) The distribution density of RMSD scores for the AF2-predicted ABL conformations using shallow MSA depth with respect to the active ABL state (orange filled bars) and the inactive states I_1_ (red filled bars) and I_2_ (blue filled bars). (C) The distribution density of RMSD scores for the AF2-predicted A-loop conformations with respect to the A-loop residues in the active ABL state (orange filled bars), the inactive states I_1_ (red filled bars) and I_2_ (blue filled bars). (D) Structural alignment of the AF2-predicted conformational ensemble obtained with AF2 and shallow MSA subsampling approach. (E) Structural overlay of the regulatory DFG motif conformations from the AF2-predicted conformational ensemble.

This analysis confirmed that the AF2 ensemble generated using shallow MSA depth adaptation can accurately predict the active ABL kinase form and capture the mobility of the A-loop seen in both the active and intermediate inactive I_1_ form. Structural alignment of the AF2-predicted structural ensemble indicated a moderate degree of conformational heterogeneity, particularly in the A-loop (Figure 6D), while conformational differences in the DFG motif are reflected in modest deviations around dominant DFG-in orientation characteristic of the active state and small displacements of the F401 residue (Figure 6E). These results are consistent with AF2 modeling of the active substrate-bound conformations for 437 catalytically competent human protein kinase domains showing that best models selected using pLDDT assessment of the A-loop residues can correctly single out the catalytically active protein kinase structure.^43^

To summarize, the analysis of the AF2 predictions using shallow MSA depth adaptation showed the reliability of predicting both the active ABL structure and conformational heterogeneity of the active ABL form. In addition, a close correspondence was found between the best predicted A-loop conformations and experimentally observed A-loop structures in the active and also intermediate I_1_ forms. However, these experiments revealed only a limited variability of the regulatory DFG-in conformation that remained confined to its active form. As a result, although subsampling of MSA and reducing MSA depth may increase conformational heterogeneity around the ground active ABL state, this approach cannot readily generate distinct low-populated inactive ABL conformations. This analysis also highlighted the limitations of this approach to capture the full heterogeneity and diversity of the ABL conformational states as the key functional regions including the αC-helix and DFG motif remained in their respective active αC-in and DF-in orientations.

### Exploring Protein Flexibility and Prediction of ABL Conformational Ensembles by SPEACH_AF Approach

In the next round of computational experiments, we explored SPEACH_AF method^14^ in which the initial AF2 models are scanned to identify interaction surfaces within the structure followed by *in silico* alanine mutagenesis of MSAs either in a systematic manner or based on possible contact points in the initial AF2 structure. We evaluated the predictive ability of this approach by generating different alanine-mutated sets of MSAs and then running shallow depth MSA pipeline to further increase structural variability. We generated multiple distinct MSAs using the SPEACH AF approach^14^ by using different random alanine mutation positions in the MSAs as well as the number of mutations in the MSAs. We then selected two MSAs out of these, with the first MSA also having a lower frequency of mutation than the second one (Figure 3). These experiments examined whether SPEACH_AF by altering the MSAs can redirect the attention of the network to distinct parts of the MSA and find ABL kinase-specific conformational ensemble patterns and predict functional conformations.

We focused our analysis and discussion on two MSAs showing a range of scenarios and predictions by SPEECH_AF adaptation (Figure 7). The analysis of pLDDT profiles for two different sets of predictions (Figure 7A,B) revealed a considerable variation in the profiles. The pLDDT values ranged from stable profile of top five models for the first SPEACH_AF experiment (Figure 8A) to extremely heterogeneous ensemble of top models derived from another SPEACH_AF alanine-mutated MSA (Figure 7B). The first pLDDT profile resembles the profile obtained from AF2 default shallow MSA setup displaying high pLDDT values for the ABL residues and showing some moderate reduction for the A-loop residues (Figure 7).

**Figure 7.**
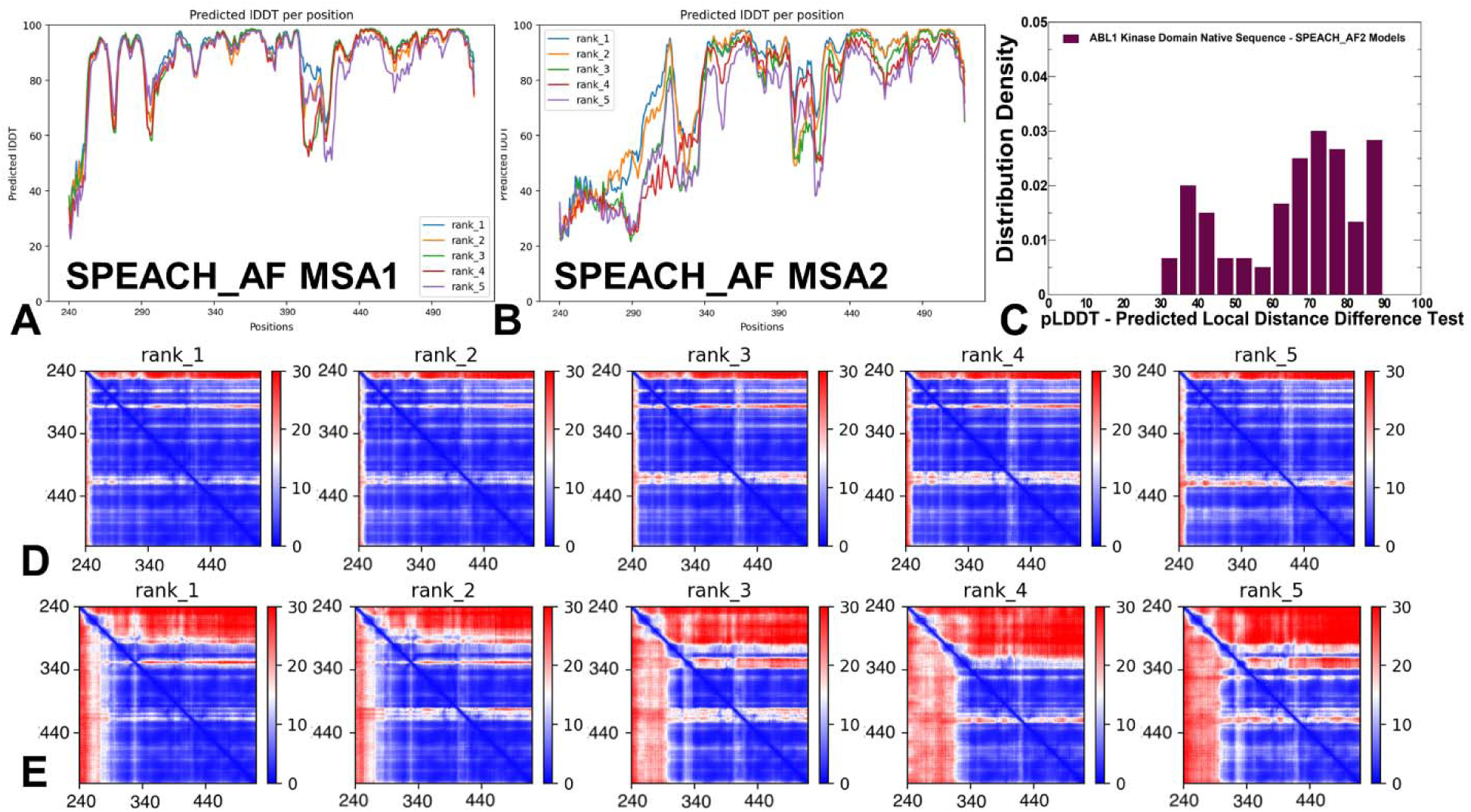
The statistical analysis of the SPEACH_AF models for the ABL kinase structures. (A,B) The residue-based pLDDT profile for the top five ranked models is obtained from two distinct sets of SPEACH_AF predictions for ABL conformations. The AF2 results on two MSAs show diversity of predicted ABL conformations by SPEACH_AF method. (C) The distributions of pLDDT structural model assessment for the ABL conformational ensemble are obtained from SPEACH_AF by using different random alanine mutation positions in the MSAs as well as the number of mutations in the MSAs. (D) The PAE heatmaps for the top five models obtained from the first SPEACH_AF set (E). The PAE heatmaps for the top five models obtained from the second SPEACH_AF set. The heat maps show the PAE between each residue in the top ranked models. The color scale contains three colors to highlight the contrast between the high confidence regions and the low confidence regions.

In sharp contrast, alanine mutagenesis of the MSA in the second experiment produced somewhat “uncontrolled” increase in conformational variability across the entire ABL kinase domain, particularly featuring low-to-moderate pLDDT values ∼40-70 for the N-terminal residues and A-loop (Figure 7B). The density distribution of the pLDDT values showed a very broadly distributed profile with one peak for pLDDT∼75-80 and another peak for pLDDT ∼40-50 where the latter peak is associated with the predictions from second SPEACH_AF experiment (Figure 7C). The heat maps of PAE values reflected these differences, showing a consistent and stable pattern for the first round of SPEACH_AF predictions (Figure 7D) and highly heterogeneous ensemble with PAEs signaling considerable level of disorder for the A-loop conformations from the second MSA predictions (Figure 7E).

The distributions of TM scores with respect to the active and two inactive conformations showed a remarkably broad range of values between 0.2 and 0.9 confirming that this approach can produce a wide range of alternative structures (Figure 8A). Interestingly, significant broad peaks at TM ∼0.6-0.9 values measuring similarities with the active form indicated that predictions continue to be dominated by active-like ABL conformations. At the same time, and unexpectedly, a strong peak of TM ∼0.85-0.9 for comparing structural similarities of the predicted conformations to the inactive I_1_ state revealed that SPEACH_AF approach can successfully recover both active and intermediate inactive conformations (Figure 8A). A more detailed quantitative analysis of the RMSDs for the complete ABL structure (Figure 8B) and A-loop (Figure 8C) revealed a broad distribution for RMSDs with respect to the active ABL conformation (RMSD∼0.4-0.8 Å) and sharp peak at RMSDs ∼ 1.0 Å -1.5 Å for A-loop. A minor and broader peak at RMSD ∼3Å-4Å compared to the A-loop of the active structure confirmed the ability of the method to also capture conformational heterogeneity of the active structure.

**Figure 8.**
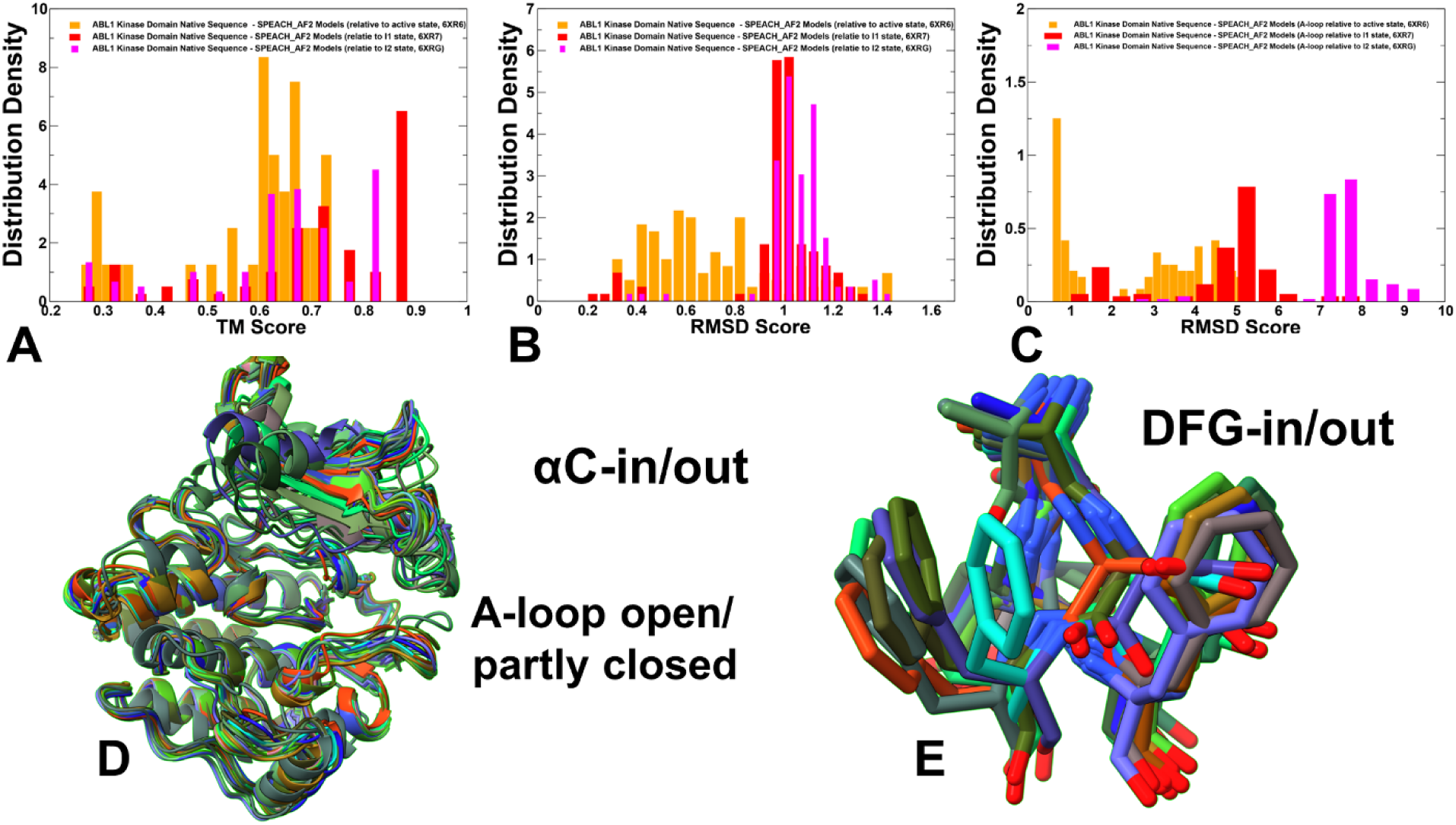
The distribution of structural similarity metrics TM scores and RMSD values for the SPEACH_AF predicted ABL structure and for predicted conformations of the A-loop with respect to the active and inactive ABL states. (A) The distribution density of TM scores for the SPEACH_AF predicted ABL conformations with respect to the experimental active ABL state (orange filled bars) and the inactive states I_1_ (red filled bars) and I_2_ (blue filled bars). (B) The distribution density of RMSD scores for the SPEACH_AF predicted ABL conformations relative to the active ABL state (orange filled bars) and the inactive states I_1_ (red filled bars) and I_2_ (blue filled bars). (C) The distribution density of RMSD scores for the SPEACH_AF predicted A-loop conformations with respect to the A-loop residues in the active ABL state (orange filled bars), the inactive states I_1_ (red filled bars) and I_2_ (blue filled bars). (D) Structural alignment of the AF2-predicted conformational ensemble obtained with SPEACH_AF approach. (E) Structural overlay of the regulatory DFG motif conformations from the SPEACH_AF predicted conformational ensemble.

The RMSD distribution for the A-loop residues displayed small peak at RMSD∼1.5Å - 2Å with respect to the inactive I_1_ structure but only relatively large RMSDs ∼ 7.0Å-8.0Å when compared to the flipped A-loop conformation in the fully inactive I_2_ (Figure 8B,C). We noticed that the increased conformational diversity of the SPEACH_AF ensembles may often yield misfolded conformations as evident from the broad span of TM scores. By examining the ensemble of SPEECH_AF conformations, we removed the misfolded and disordered conformations with low pLDDT values and present the alignment of only functionally relevant ABL conformations (Figure 8D) featuring significant displacements of the functional kinase regions P-loop, αC-helix and A-loop. Notably, the diversity of the predicted conformations is also reflected in the distribution of DFG conformations showing markedly increased variability and extensive sampling of multiple intermediate DFG-out positions (Figure 8E). Consistent with the experimental analysis of the inactive I_1_ state, SPEACH_AF predicted conformational ensemble featured the DFG motif flipped from active DFG-in position by 180° and adopting the inactive DFG-out conformation. At the same time, the vast majority of the SPEACH_AF generated conformations featured the A-loop in the open conformation similar to the active form (Figure 8E). Because of the DFG flip, the neighboring residues of the DFG motif (L403-M407) are also flipped by 180°. We also noticed that D400 of the DFG motif is in the “out*”* conformation and can swap positions with F401 (Figure 8E).

Hence, our results demonstrated that SPEACH_AF adaptation of AF2 can enhance conformational heterogeneity of the predicted kinase states but could often produce misfolded and partially unfolded conformations. The premise of this approach is that if alanine mutations alter the information content of the MSA, the AF2 models are expected to produce a multitude of alternate conformations. According to our results, this method can only partly capture conformational heterogeneity of the thermodynamically dominant active structure while also predicting structural transformations in the A-loop and DFG motif seen in the inactive I_1_ state. However, despite markedly increased variability of the AF2-predicted ABL conformations, this approach could not detect the functionally important inactive I_2_ structure which is dramatically different from both active state and inactive state I_1_ . In the I_2_ structure, the A-loop rotates with the middle of the A-loop translating by ∼ 35 Å while F401 of the DFG motif not only flips into the catalytic pocket as was observed in I_1_ but undergoes a massive conformational swing by ∼11 Å to occupy a hydrophobic pocket that blocks binding of ATP or competitive inhibitors.^38^ Structural changes associated with the transition to the I_2_ structure are large and are executed through coordinated massive rearrangements of the key structural elements P-loop, A-loop, and αC-helix. Whereas the DFG motif is in the “out*”* conformation in both I_1_ and I_2_ states, the A-loop and the αC helix adopt vastly different conformations in these states.

### Randomized Alanine Sequence Scanning and Shallow MSA Subsampling Facilitate AF2 Prediction of Structural Ensembles for the Conformational States of ABL1 Kinase

We introduce a randomized alanine scanning adaptation of the AF2 methodology in which the algorithm operates first on the pool of sequences and iterates through each amino acid in the native sequence to randomly substitute 5-15% of the residues with alanine, thus emulating random alanine mutagenesis. The algorithm substitutes residue a_i_ with alanine at each position i with a probability p_i_ randomly generated between 0.05 and 0.15 for each sequence position. Specifically for the proposed protocol, randomized alanine sequence scanning is followed by construction of corresponding MSAs and then by AF2 shallow subsampling applied on each of these MSAs. Using this approach (see Materials and Methods for more details) we predicted structures and conformational ensembles of the ABL kinase. The central distinguishing feature of the proposed AF2 adaptation is that the proposed algorithm can moderately and systematically perturb the MSAs, while avoiding introduction of random mutations in the homologous sequences within MSAs themselves, which is also the major difference from the SPEACH_AF approach (Figure 9).

The analysis of pLDDT distribution densities for produced conformations revealed an expected and generally conserved pattern^57^ in which we observed high confidence pLDDT values are consistently found for the ABL kinase domain in the thermodynamically dominant active state (Figure 9A-C) while the lowered pLDDT values are featured for the A-loop (residues 398-421) featuring both an active DFG-in and inactive DFG-out state (Figure 10A-C). Another key regulatory element of the ABL kinase is the αC-helix (residues 292-312) where αC-helix-in position is characteristic of the active state (Figure 9) while αC-helix-out position is present in intermediate and inactive kinase forms (Figure 10).

For some AF2 experiments using randomized alanine sequence scanning (Figure 9), the AF2 predictions produced consistently high pLDDT values for the functional kinase regions, including A-loop and αC-helix, which typically reflected accurate predictions of the active ABL state. In addition, the AF2 predictions correctly reported the reduced confidence in the A-loop residues as the A-loop conformation cannot be adequately represented by a single structure. We also observed a considerable variability among the pLDDT distributions, particularly in the N-terminal regions, C-terminal loops and especially in the A-loop (Figure 9). For some other randomized alanine scanning AF2 experiments (Figure 10) we observed the reduced pLDDT values for the A-loop and considerable variation in the A-loop prediction assessment among the top five models. The relatively moderate pLDDT values were observed for the αC-helix residues and we found ABL conformations with both active αC-in and inactive αC-out orientations.

**Figure 9.**
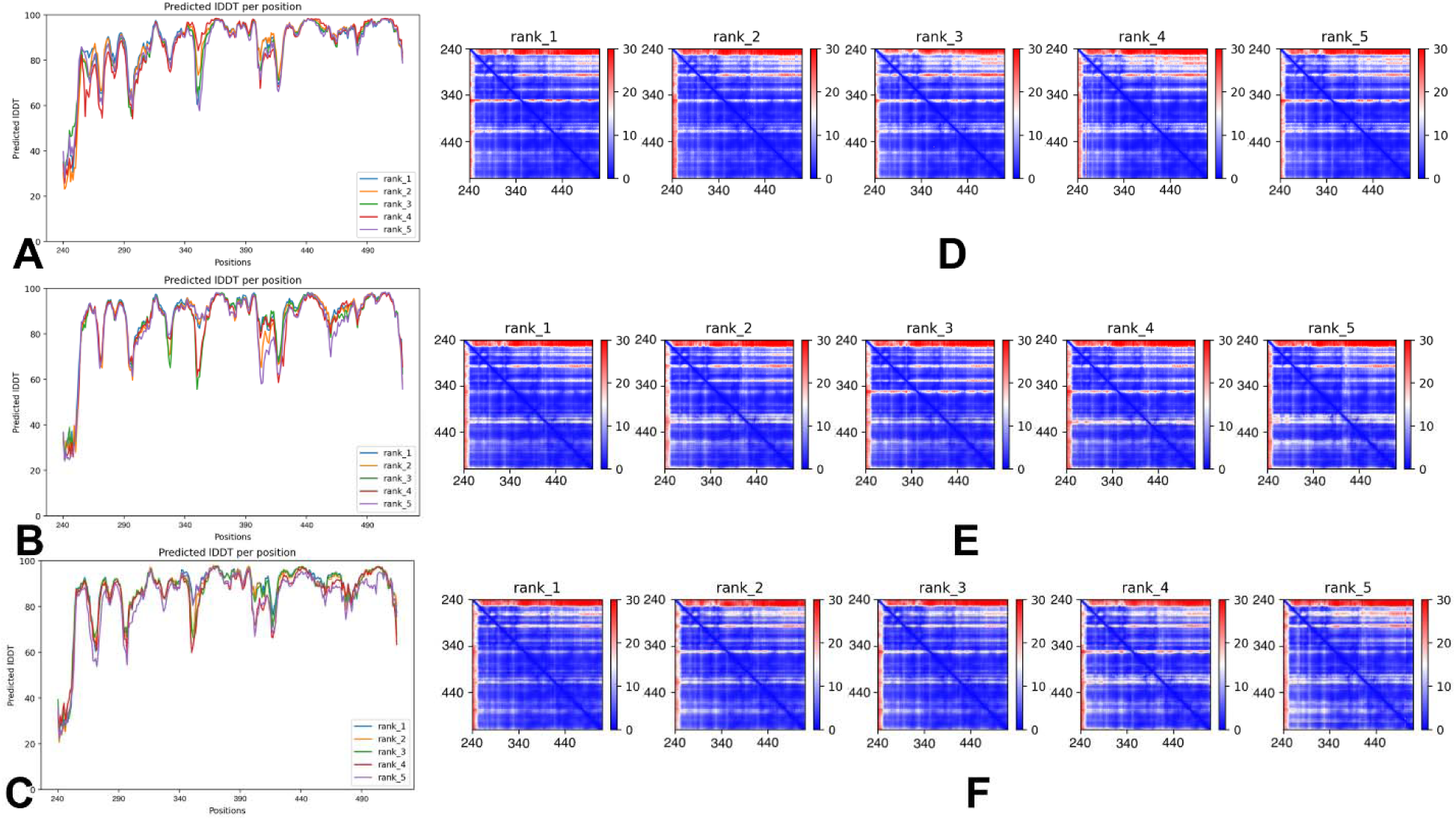
The statistical analysis of the predicted AF2 models for the ABL kinase structures using randomized alanine sequence scanning adaptation of AF2. We report the results of the approach obtained from using the algorithm multiple times on the full native sequence, resulting in distinct sequences, each with different frequency and position of alanine mutations. MSAs were then constructed for each of the mutated sequences using the alanine-scanned full-length sequences as input for the MMSeqs2 program followed by shallow MSA AF2-based structure generation. The panels (A)-(C) depict the produced residue-based pLDDT profiles of the top five ranked models in three different randomized sequence scanning/AF2 experiments. These experiments highlight cases of consistent predictions of the active ABL state with high confidence pLDDT values for the protein residues. The high pLDDT values and the patterns of PAE profiles highlight consistent and accurate prediction of the A-loop conformation (residues 398-421) and other functional regions for the active ABL state.

These experiments highlighted the diversity of the predicted inactive ABL conformations demonstrating a pronounced variability for the A-loop (residues 398-421) and other functional regions. The PAE heatmaps between each residue in the model (Figure 10) underscored the differences between the high confidence regions and the low confidence regions, particularly demonstrating the increased heterogeneity of the A-loop and C-lobe residues in the predicted models. These findings suggested that the randomized alanine sequence scanning adaptation of the AF2 approach can produce a broad ensemble of both active and inactive I_1_ states. The results also indicated that the A-loop and C-lobe regions become increasingly flexible in the intermediate inactive forms which is consistent with the kinase regulation mechanisms where these functional regions undergo “cracking” to facilitate considerable structural changes during transitions between the inactive and active kinase states.^58^ In general, the pLDDT distributions obtained using this approach highlighted conformational diversity of the AF2-predicted states while maintaining a “controllable” level of variability in the predicted ABL ensemble, avoiding any misfolded predictions and capturing functional conformational heterogeneity among both active and inactive states (Figure 9). This should be contrasted to what we found to be a more “radical” variations in flexibility when using SPEACH_AF approach, which often produced excessive mobility of the ABL core as well as often emerging misfolded structures.

The cumulative density distribution of the pLDDT values (Figure 11A) displayed a strong peak at pLDDT∼85-90 and exhibited several more peaks at lower pLDDT values that become smaller as pLDDT lowers (Figure 11A). The broad coverage of TM score values for the predicted conformations coexists with peaks at TM score ∼0.8-0.9 showing structural similarities with the active and inactive I_1_ conformations (Figure 11B).

**Figure 10.**
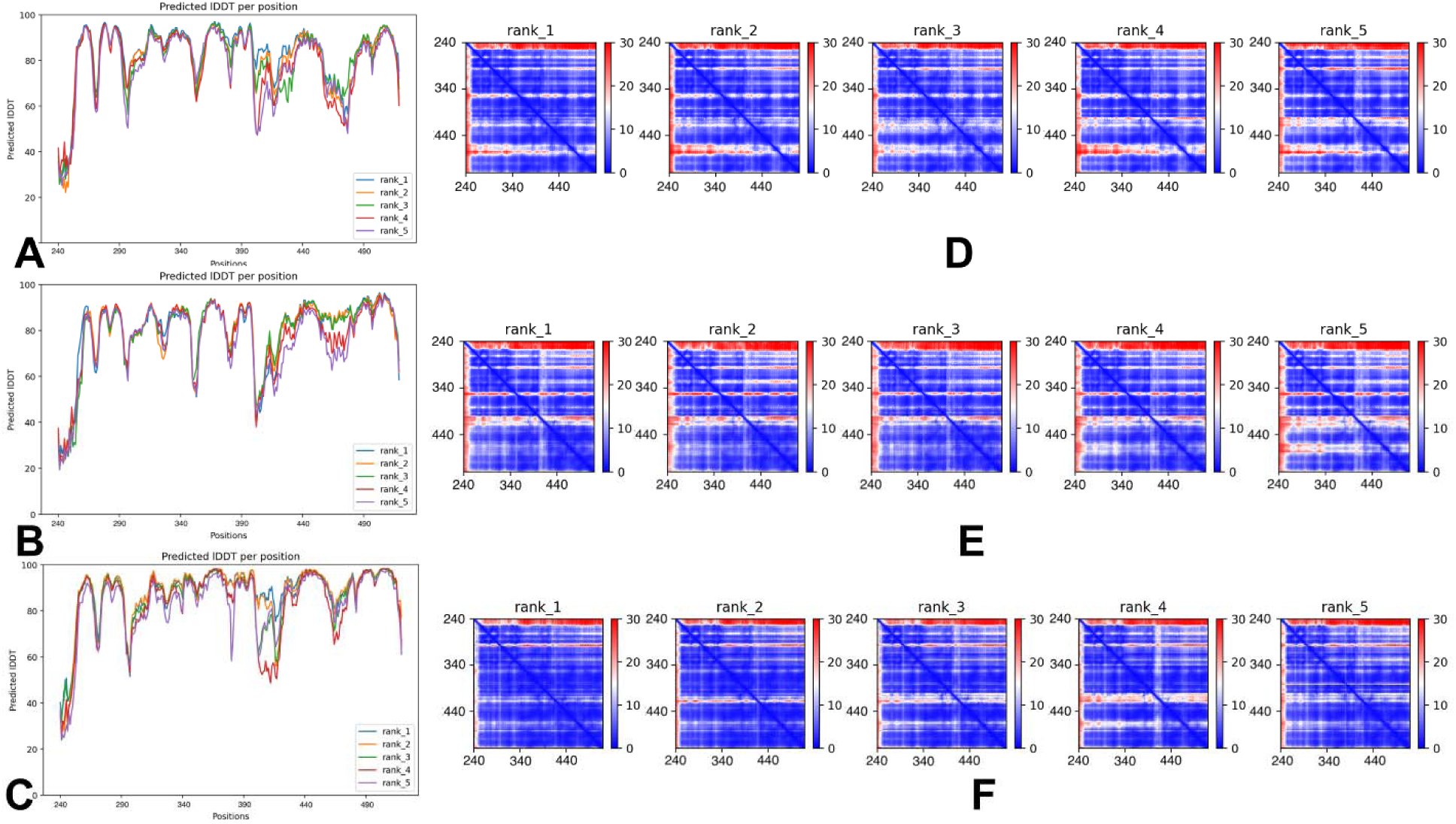
The statistical analysis of the predicted AF2 models for the ABL kinase structures using randomized alanine sequence scanning adaptation of AF2. The panels (A)-(C) depict the produced residue-based pLDDT profiles of the top five ranked models in three different randomized sequence scanning/AF2 experiments. These experiments highlight the diversity of the predicted ABL conformations (active and inactive) where top ranked models can differ appreciably in their pLDDT values. The pLDDT and PAE profiles highlight a pronounced variability for the A-loop (residues 398-421) and other functional regions.

The distribution of the RMSD values for the A-loop residues showed distinct peaks showing that most of the predicted conformations correspond to the ensemble of active conformations (RMSD < 1.0 Å) (Figure 11C,D). There is another peak at RMSD ∼ 3.0-3.5 Å from the inactive I_1_ structure indicating that a significant population of predicted conformations are structurally similar to the inactive I_1_ structure. Notably, the RMSD distribution of predicted conformations with respect to the inactive I_2_ conformation showed a peak at larger RMSD values and a small but noticeable population of the predicted ABL conformations with the A-loop residues within RMSD ∼ 3.0 Å from the inactive I_2_ state (Figure 11D). Although our predictions of the conformational ensembles cannot be directly compared with the NMR experiments showing that the relative state population of the ABL1 ground active state in solution is 88% and populations of I_1_ and I_2_ are ∼ 6%, we found that the active conformation is the most dominant ABL state in the AF2-predicted ensemble. Importantly, combining randomized alanine scanning with MSA construction and AF2 predictions with shallow MSAs also produced a significant functionally relevant ensemble of the heterogeneous inactive I_1_ conformations, while only a small fraction of the generated conformations are similar to the fully inactive I_2_ structure with the flipped A-loop and inactive DFG-out motif (Figure 11).

**Figure 11.**
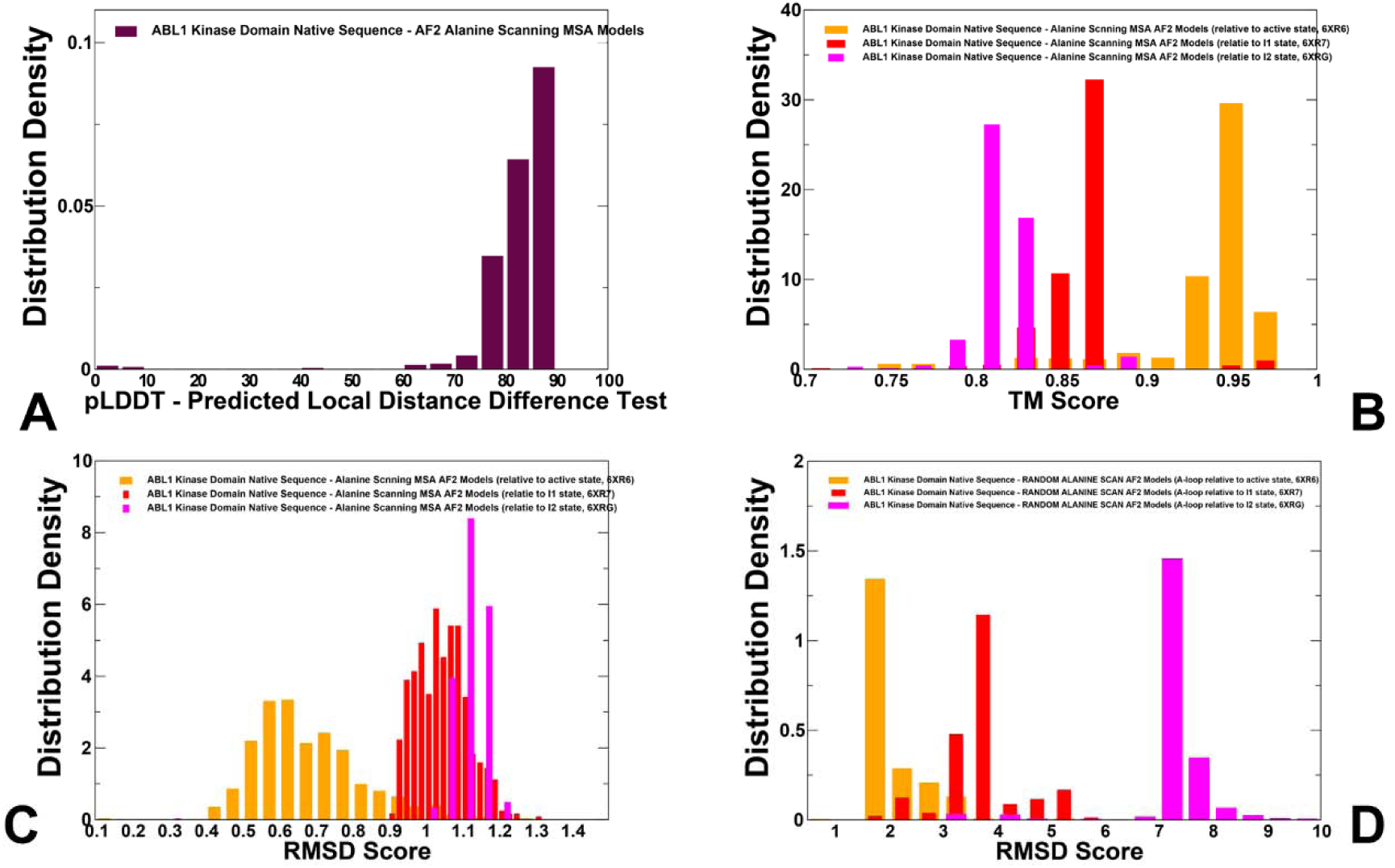
The analysis of AF2 predictions using randomized alanine sequence scanning approach. The distribution of pLDDT assessment metric and structural similarity metrics TM scores and RMSD values for the AF2 predicted structure and for the predicted conformations of the A-loop with respect to the active and inactive ABL states. (A) The cumulative residue-based pLDDT profile for the top five ranked models is obtained from nine different experiments using AF2 with randomized alanine scanning approach. (B) The distribution density of TM scores for the AF2-predicted ABL conformations with respect to the experimental active ABL state (orange filled bars) and the inactive states I_1_ (red filled bars) and I_2_ (blue filled bars). (B) The distribution density of RMSD scores for the AF2-predicted ABL conformations relative to the active ABL state (orange filled bars) and the inactive states I_1_ (red filled bars) and I_2_ (blue filled bars). (C) The distribution density of RMSD scores for the AF2-predicted A-loop conformations with respect to the A-loop residues in the active ABL state (orange filled bars), the inactive states I_1_ (red filled bars) and I_2_ (blue filled bars).

Structural mapping of the AF2-predicted conformational ensemble demonstrated a functionally significant variability of the ABL kinase domain, suggesting that randomized alanine scanning approach combined with shallow MSA AF2 could produce conformational ensemble reflecting functional heterogeneity of the kinase domain that is dominated by the active ABL conformation with the αC-in, DFG-in, and a highly flexible A-loop in its open conformational form (Figure 12A). By projecting the complete range of AF2-produced DFG conformations, we illustrated the functionally relevant positional variability of the DFG motif and a “continuous” spectrum of movements between active DFG-in position and inactive DFG-out conformations (Figure 12B). The variability of the DFG motif is particularly exemplified by the observed movements of F401 residue that samples a large number of intermediate states between DFG-in and DFG-out flipped by 180° (Figure 12B). In addition to the active ABL state, the AF2 ensemble generated a significant population of ABL inactive I_1_ conformations that sample changes between αC-in and αC-out inactive positions as well as intermediate DFG-in to DFG-out orientations (Figure 12).

**Figure 12.**
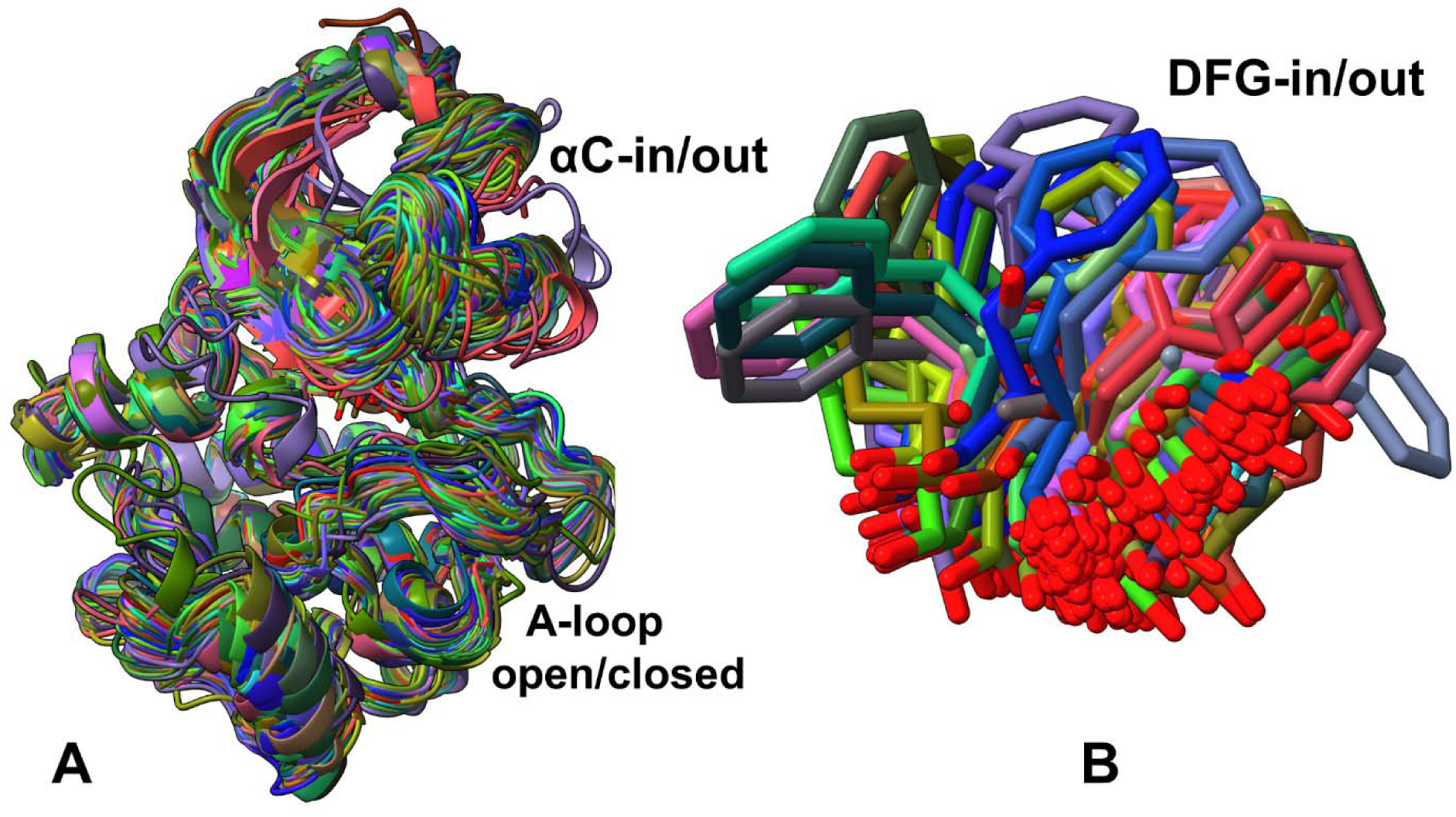
(A) Structural alignment of the AF2-predicted conformational ensemble using randomized alanine sequence scanning. (B) Structural overlay of the regulatory DFG motif conformations from the AF2-predicted conformational ensemble.

It is worth noting that structural alignment of the DFG conformations in the I_1_ NMR ensemble showed that DFG motif is flipped from the active “in” position and can adopt a spectrum of “out” conformations while A-loop remains flexible and in the open active form (Figure 1E). The diversity of the predicted ABL conformations is illustrated by selecting AF2 models that are close to the inactive I_1_ state (RMSD < 1.5 Å) (Supplementary Information, Figure S2). Strikingly, the kinase core and the A-loop conformation shared by these models are similar and closely resemble the I_1_ state, but the DFG motif can adopt a number of intermediate DFG-out positions (Figure 12, Supplementary Information, Figure S2). In addition, most of the AF2-predicted ABL states tend to shift αC helix towards the inactive αC-out orientation (Supplementary Information, Figure S2A).

To understand functional relevance of the predicted inactive conformations, the AF2 ensemble was examined using the recently proposed nomenclature for the structures of active and inactive kinases.^46^ Interestingly, a significant fraction of the predicted inactive conformations directly correspond to the “BLBplus “ class (DFG-in/out, αC-helix-out) that represents one of the two common inactive kinase forms^46^ where αC-helix assumes inactive αC-out conformation and DFG-Phe motif samples intermediate “out” positions with F401pointing upward and thus pushing the αC-helix outward (Figure 12, Supplementary Information, Figure S2B). According to the statistical analysis of the active and inactive kinase forms, 95% of kinase structures with the active DFG-in conformation also have the active αC-in position, while 77% of kinase structures with the inactive DFG-in/out upward conformation have the αC-helix out conformation In this conformation, the DFG-Phe ring is underneath the C-helix but pointing upward forcing the C-helix outwards.^46^ Hence, the important finding of this analysis is that the proposed random alanine sequence scanning adaptation combined with shallow MSA AF2 modeling enables prediction of functionally significant conformational heterogeneity in the ABL ensemble, capturing the dominant population of the active ABL form along with the populations of largely heterogeneous I_1_ ensemble of conformations.

The presented results can be better interpreted and understood in the context of the experimental structural studies and our previous atomistic simulations of the ABL kinase states. The results of our previous MD simulations and MSM analysis^42^ suggested that the ABL kinase domain can utilize the inactive state I_1_ for pathways between the active form and the inactive state I_2_. Consistent with simulation data, the results of the AF2 predictions indicated that a broad ensemble of inactive intermediate BLBplus conformations (DFG-in/out, C-helix-out) may effectively connect the active form with the inactive state I_1._ (Figure 12). Importantly, our AF2 prediction results are also consistent with the recent unbiased MD simulations of the ABL-Imatinib binding, capturing conformational change from the DFG-in/A-loop open conformation to the DFG-out/A-loop closed conformation.^59^ This study showed that binding involved four primary conformational states—the DFG-in/A-loop open and DFG-out/A-loop open states of the apo kinase, as well as the DFG-out/A-loop intermediate state. At the same time, the fully inactive DFG-out/A-loop closed form is formed and becomes more stable only in the Imatinib-bound kinase.^59^ Consistent with this study, the AF2-predicted conformations are associated with predominantly active DFG-in/A-loop open conformation and the inactive conformation, where the αC-helix assumes inactive “out” orientation, DFG motif is “flipped” to adopt a “DFG-out” conformation, while the A-loop remains to be in the open, active-like form (Figure 12). According to the unbiased MD simulations of the ABL-Imatinib binding, the inhibitor first selectively recognizes the flipped DFG-out/A-loop open kinase conformation followed by entering the ATP-binding site after which the inhibitor induces the closure of the A-loop in the fully inactive state.^59^ Our results are fully consistent with these pioneering computational studies, showing that applying randomized alanine sequence scanning together with AF2 shallow MSA framework generates accurate conformational ensembles of the active and intermediate inactive states (αC-out, DFG-out, A-loop open) that play a central role in the ABL-drug recognition and binding process. Our results showing that the prediction of population of the fully inactive I_2_ state (DFG-out, αC-helix-out and closed A-loop) are rare also echoed in observations that this ABL conformation is transient and “hidden” on the unbound ABL landscape and it may become stable only upon Imatinib binding.^59^ Our results reach similar conclusions as recent illuminating analysis of fold-switching proteins, showing that AF2 predictions of these allosteric systems are strongly biased towards predictions of the more stable ground fold-switch state while often failing to detect less stable “excited” stats.^16,17,60^

### Local Frustration Analysis of the ABL Structures and AF2-Structural Ensembles

Here, we propose a physical model for analyzing and interpreting the results of AF2 predictions of ABL allosteric states by hypothesizing that the accurate AF2 predictions of structural ensembles of the active and intermediate I_1_ states and inability to consistently locate the inactive I_2_ form may be associated with the radical differences in local frustration patterns of these states. We examine local frustration patterns of the ABL states and show that the AF2-predicted thermodynamically dominant active and inactive I_1_ states are characterized by minimally frustrated conformational landscapes that ensure both stability and kinetic accessibility of these conformations. On the other hand, we will demonstrate that the fully inactive I_2_ state is highly frustrated in the critical functional regions A-loop, αC-helix forming non-native and energetically unfavorable interactions in segments involved in the active-inactive switching. Our hypothesis for rationalization of the AF2 predictions is based on the notion that the minimally frustrated interactions of thermodynamically dominant states within a protein family can be conserved over evolutionary timescales which implies their driving role in determining the foldability and coevolutionary signals which are learned and then captured by the AF2 methodology. We propose that the AF2 inability to accurately predict all multiple experimental conformations of regulatory switchers such as ABL kinase may be also associated with the intrinsic differences in local frustration of conformational landscapes for the thermodynamically favorable and low-populated excited states. In this model, the highly frustrated, non-native interactions in alternative excited kinase states can be associated with specific coevolutionary signals relevant for alternative states that are not learned by the AF2 architecture which forces AF2 predictions with different randomized adaptations towards minimally frustrated and thermodynamically favorable states. The analysis is based on the profiling of the ABL residues in different states by local frustratometer^50–52^ which computes the local energetic frustration index using the contribution of a residue to the energy in a given conformation as compared to the energies that would be found by mutating residues in the same native location or by creating by changing local conformational environment for the interacting pair.

The conformational frustration profiles showed moderate high frustration density for the active state (Figure 13A) and inactive state I_1_ (Figure 13B) with the A-loop in the fully extended or open states. Importantly, it can be noticed that the high frustration density for the A-loop (residues 398-421) is fairly small in both the active and I_1_ states as the corresponding density profiles for the A-loop residues displayed local minima (Figure 13A,B) while the minimally frustrated density profile showed moderate peaks for the A-loop positions (Figure 13D,E). Interestingly, the shape of the local frustration density profiles for the active and the inactive I_1_ states looked similar, with some exceptions associated with the high frustration for the P-loop region (residues 260-280) in the I_1_ form (Figure 13B). The DFG motif is flipped 180° in the I_1_ form with respect to the active conformation, and it adopts the DFG-out conformation. However, the A-loop remains in an open conformation similar to the active conformation. The frustration profiles near DFG motif and neighboring A-loop residues featured a considerable level of minimal frustration contacts which allows for moderate conformational variations without compromising the energetics of these conformations (Figure 13A,B). In contrast for the fully inactive I_2_ form, the density of highly frustrated contacts is significantly increased (Figure 13C). In particular, for the I_2_ state, the high frustration density profile revealed pronounced peaks associated with the αC-helix (residues 300-311), A-loop (402-421), the adjacent region in the C-lobe (residues 420-440) that includes the P+1 motif and the αG-helix region (residues 460-480) (Figure 13C). The P+1 segment is critical for substrate recognition, and also serves as hydrophobic glue holding the sub-domains of the C-lobe together. The APE motif (residues 426-428) is anchored to the αF, αG and αI-helices linking the activation segment and C-terminal subdomains.

**Figure 13.**
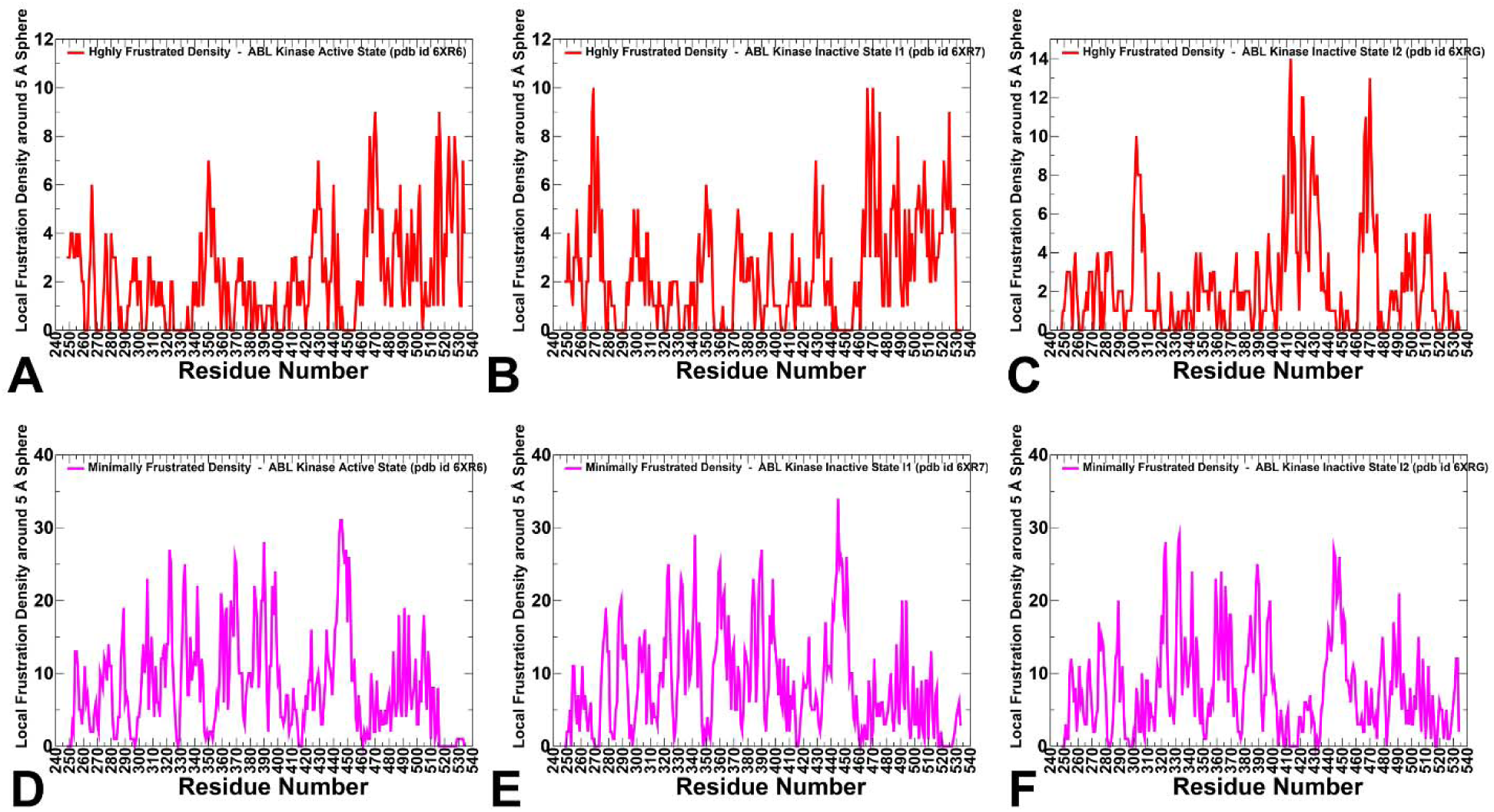
The local density of contacts distributions of conformational frustration in the ABL structures. The residue-based high frustration density in the active ABL structure (A), inactive I_1_ state (B) and inactive I_2_ state (C). The high frustration density is shown in red lines. The residue-based minimal frustration density in the active ABL structure (D), inactive I_1_ state (E) and inactive I_2_ state (F).

This highly frustrated region participates in substrate binding through the P+1 loop, and the APE-αF loop is often involved in protein–protein interactions with regulatory proteins. This can be contrasted to the active and inactive I_1_ structures in which the contacts formed by residues proximal to A-loop positions are mostly minimally frustrated (Figure 13D,E). The observed change in the local conformational frustration of the inactive I_2_ form echoed our previous studies of local frustration in the protein kinases showing that conserved locally frustrated clusters could overlap with the kinase segments involved in conformational changes associated with the kinase function.^61^ The previous large scale analysis of local frustration in protein kinases also showed that the majority of locally frustrated sites reside in the C-lobe, populating the substrate binding region of the catalytic core framed by the αF, αG, helices, and the A-loop.^61^

The conformational frustration profiles for AF2-predicted intermediate inactive states showed moderate high frustration density (Figure 14) that is reminiscent of the inactive I_1_ and also partly I_2_ state, exhibiting several specific peaks. The corresponding high frustration clusters correspond to the αC-helix (residues 300-311), the C-lobe regions including the P+1 motif (residues 420-440) and the αG-helix (residues 460-480) (Figure 14A-C). Importantly, however, these distributions revealed only moderate high frustration for the A-loop residues following DFG motif (residues 403-421), which is more reminiscent of the frustration profile of the inactive I_1_ structure but vastly different from high frustration constellation of A-loop residues see in the fully inactive, A-loop closed state (Figure 14A-C). Hence, although the AF2-predicted ensembles can accurately reproduce structures of the active and intermediate inactive states, prediction of highly frustrated A-loop region in the I_2_ state appeared to be difficult using all examined AF2 adaptations.

**Figure 14.**
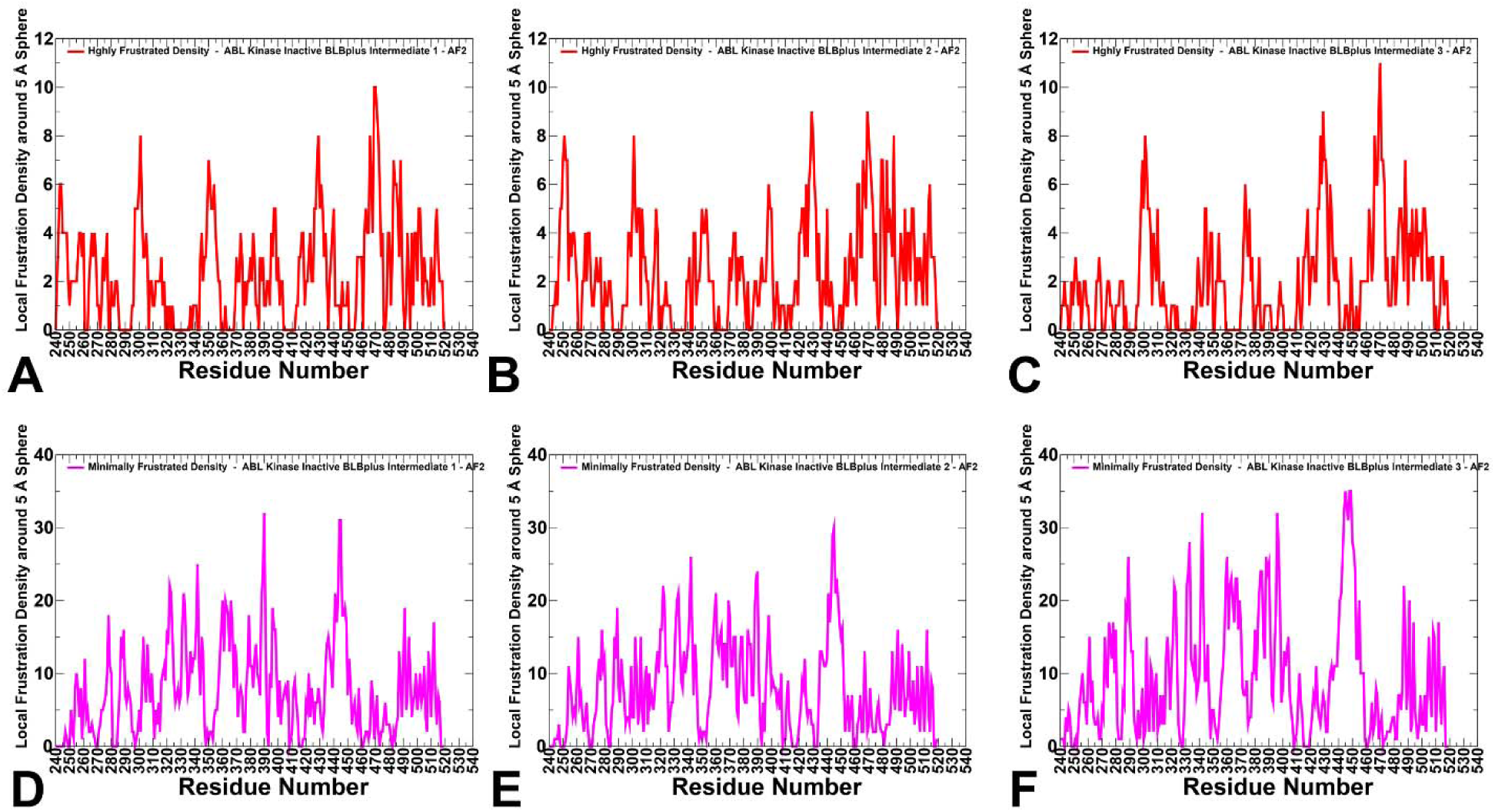
The local density of contacts distributions of conformational frustration in the ABL structures in the AF2-generated intermediate inactive structures that belong BLBplus class (DFG-in/out, αC-helix-out) of inactive conformations. The residue-based high frustration density in the three different intermediate inactive structures (A-C). The high frustration density is shown in red lines. The residue-based minimal frustration density in the three different intermediate inactive structures that belong BLBplus class (DFG-in/out, αC-helix-out) of inactive conformations (D-F).

In addition, we also performed both complete conformational and mutational frustration analysis for the ABL states and highlighted the highly frustrated, neutrally frustrated and minimally frustrated local densities (Supplementary Information, Figure S3). The results of conformational frustration analysis (Supplementary Information, Figure S3A-C) highlighted the prevalence of the neutral frustration which is consistent with previous studies of frustration in proteins showing that protein residues tend display neutral frustration which implies a moderate degree of stabilization/destabilization for the native interactions as compared to the average energetics induced by conformational or mutational changes in the respective positions.^55,56^

The local frustration analysis of the ABL states is consistent with a more general frustration analysis of different proteins that revealed the majority of local interactions as neutral (∼50-60%) or minimally frustrated ( 30%) with only 10% of the total contacts considered as highly frustrated.^50^ A direct comparison of high frustration and minimal frustration local densities is particularly revealing, showing the dominance of minimal frustration densities for the active state (Supplementary Information, Figure S3A,D). Interestingly, the profiles for the high and minimal frustration densities in the I_1_ state (Supplementary Information, Figure S3B,E) are quite similar to the active form, suggesting that both states are characterized by overall minimally frustrated landscape. In contrast, conformational and mutational frustration densities for the I_2_ state displayed high frustration density peaks in the A-loop and other functional regions (Supplementary Information, Figure S3C,F).

### Interconnected High Frustration Clusters in the Low-Populated Inactive ABL State Define Cracking Sites of Allosteric Changes and Present Difficult Targets for AF2 Assessment

To provide structure-based analysis and interpretation of local frustration patterns in ABL kinase, we mapped the top 10% of highly frustrated sites and minimally frustrated sites in the active state (Supplementary Information, Figure S4), inactive I_1_ state (Supplementary Information, Figure S5), and inactive I_2_ state (Supplementary Information, Figure S6). The high frustration sites in the active state are mostly localized in the αG-helix region (residues 460-480) with several additional isolated sites in the C-lobe (Supplementary Information, Figure S4A). Notably, the functional αC-helix and A-loop regions are minimally frustrated in the active form. The high frustration density sites become more densely populated in the inactive I_1_ state (Supplementary Information, Figure S5A) where, in addition to the C-lobe αG-helix, some of the sites are located near the DFG and αC-helix undergoing in-out shifts. However, there is no high frustration density in the A-loop that remains in the open form. Strikingly, a vastly different distribution of high frustration density sites was observed in the inactive I_2_ state (Supplementary Information, Figure S6A). We found a considerable density of high frustration sites in the αC-helix and in the C-lobe including portion of the P+1 loop (residues 420-440) and the αG-helix (residues 460-480). Notably and differently from other states, we detected a significant density of high frustration positions in the closed A-loop, mostly in the flipped segment of the A-loop (Supplementary Information, Figure S6A). Importantly, the highly frustrated sites from the A-loop, P+1 substrate binding motif and the αG-helix are clustered together forming a large fraction of C-lobe. Interestingly, the high frustration residues occupy critical regions of the inactive I_2_ state and orchestrate conformational switches between the inactive and active states. Hence, our results indicated that the ABL kinase regions undergoing large structural changes during inactive-active transitions could be enriched in clusters of highly frustrated residues. These findings are consistent with recent studies showing that identification of conserved highly frustrated contacts can determine residues involved in conformational transitions.^56^

The emergence of local frustration in these regions could also indicate the “initiation cracking points” of the inactive kinase form, ultimately facilitating global conformational transitions and shifting a dynamic equilibrium between kinase states towards the active form.^58^ Local cracking model in protein kinases is often referred to formation of unfavorable residue-residue interactions that can be compensated by increased local entropy in intermediate states during large conformational changes between inactive and active states.^58,62^ In this model, the formation of frustrated contacts and high strain may result in partial local unfolding (cracking) to facilitate large conformational transitions. Our local frustration analysis particularly indicated that a large cluster of high frustration sites in the inactive I_2_ state may be involved in such cracking mechanism during transitions to the active form. The emergence of highly frustrated clusters in the αC-helix, A-loop and C-lobe regions could present the “initiation cracking points” that could perturb the inactive state and promote dynamic functional transitions to the catalytically competent active kinase form.

Strikingly, AF2 adaptations often fail to correctly predict structural arrangement of this highly frustrated region in the I_2_ state and thus cannot accurately reconstruct the unique structure of the fully inactive form. The inactive I_2_ state which is characterized by a significant amount of local frustration and presence of large structural clusters of frustrated unfavorable contacts can feature coevolutionary signals that are masked in AF2 predictions. We argue that AF2 adaptations can predict distinct conformations that are associated with stable interactions and coevolutionary signals produced by minimally frustration contacts observed in the native protein structures. This analysis supported the emerging notion that AF2 models cannot readily predict conformational mechanisms driven by frustration changes and allosteric transformations. From the point of view of localized frustration, proteins maintain a conserved network of minimally frustrated interactions in the thermodynamically dominant state. Although a broad web of minimally frustrated residues in the kinase catalytic core could reflect robustness of the protein kinase fold to evolutionary pressure and mutations, the presence of locally frustrated residue clusters may be required for tailoring protein kinase dynamics to maintain a dynamic equilibrium between alternative kinase states required for normal function. The results of this study suggest that AF2 prediction pipelines are biased towards robust predictions of minimally frustrated, thermodynamically favorable states as coevolutionary signals inferred from these structures dominate AF2 training and inference networks. The significant algorithmic dependence of AI structure prediction algorithms on coevolutionary informational signals explains the lack of predicted power for the rare inactive ABL kinase state which may be determined by the fact that this functional conformation represents transient evolutionary byproduct with significant frustration content (both geometrical and energetic) that are not readily discernible by learning of coevolutionary signals in protein families.

## Conclusions

In the current study, we employ and systematically compare several recent AF2 adaptations including (a) MSA subsampling with shallow MSA depth and (b) SPEACH_AF approach in which alanine scanning is performed on generated MSAs by using different random alanine mutation positions in the MSAs. In addition, we introduce a random alanine scanning algorithm that iterates through each amino acid in the native sequence and randomly substitutes 5-15% of the residues to simulate random alanine substitution mutations. For each of the mutated sequences MSAs are constructed that are then enter the AF2 prediction pipeline. By systematically applying and comparing these three AF2 pipelines along with probing of the AF2 parameters, we predict the structures and populations of the ABL conformational ensembles. We show that MSA shallow subsampling approach can adequately predict the active ABL conformation and the conformational heterogeneity around the active form but is unable to reliably reproduce the low-populated inactive conformations. A comparative analysis reveals that SPEACH_AF approach can increase the conformational diversity by sampling both open and partly closed A-loop conformations but may be sensitive to the mutation-generated MSAs and frequency of mutations in the MSAs often producing highly disordered and misfolded conformations. We show that the randomized alanine scanning of the native sequences followed by construction of shallow MSAs, and AF2-based structure generation allows for controllable perturbation of MSAs without mutating the homologous sequences within them, which is the major difference from the SPEACH_AF approach. We suggest that this approach can allow for gradual diversification of the attention network mechanism and discover distinct patterns of coevolved residues from the MSAs. The results demonstrate that the proposed AF2 adaptation enables accurate and robust prediction of structural ensembles and conformational heterogeneity of the active and the intermediate inactive ABL state.

By employing local frustration analysis of the AF2-predicted conformations, this study unveiled previously unappreciated connections between local frustration patterns of the kinase conformational states and the ability of the AF2 methods to predict structural ensembles of the active and inactive states. We show that the dominant minimal frustration pattern in the active ABL state and the inactive intermediate I_1_ state provide a broad and funnel-liked landscapes around these states allowing AF2 predictions to accurately capture structural ensembles of these functional ABL conformations. In contrast, the emergence of interconnected high frustration residue clusters in the inactive I_2_ state that define the initiation “cracking” sites of allosteric changes and present difficult targets for robust AF2 predictions. We argue that since AF2 relies on the training set of minimally frustrated, thermodynamically stable protein states, coevolutionary patterns of MSAs may guide and bias prediction of structural ensembles towards dominant kinase states. At the same time, highly frustrated inactive states are rare for sufficient training and robust inference of these states in the predicted conformational ensembles. The results of this study revealed both the power and intrinsic limitations of the AF2-based methods and new adaptations, showing that multi-fold coevolutionary signals may arise from the sequences of protein subfamilies characterized by diverse conformations and that incorporation of local frustration information and attention-based learning of both minimal and high frustration patterns across protein folds may augment the predictive abilities of the AF2-inspired methods.

## Data Availability Statement

Data is fully contained within the article and Supplementary Information material. Crystal structures were obtained and downloaded from the Protein Data Bank (http://www.rcsb.org). The rendering of protein structures was done with UCSF ChimeraX package (https://www.rbvi.ucsf.edu/chimerax/) and Pymol (https://pymol.org/2/) . The software tools used in this study are freely available at GitHub sites :

https://github.com/deepmind/alphafold; https://github.com/sokrypton/ColabFold/; https://github.com/RSvan/SPEACH_AF; https://www.github.com/HWaymentSteele/AFCluster; https://github.com/smu-tao-group/protein-VAE.

All the data obtained in this work, the software tools, and the in-house scripts are freely available at ZENODO general-purpose open repository: https://zenodo.org/records/10656097.

## Author Contributions

Conceptualization, G.V.; Methodology, N.R., M.A., G.C., H.T., S.X., G.V. P.T.; Software, N.R., H.T., S.X., M.A., G.G., P.T., G.V. Validation, N.R., G.V.; Formal analysis, N.R., G.V., M.A., G.G., H.T., S.X., and P.T.; Investigation, N.R., G.V. and P.T.; Resources, N.R., G.V., M.A, H.T., S.X., and G.V. Data curation, N.R., M.A., G.C., H.T., S.X., G.V. Writing—original draft preparation, N.R., G.V.; Writing—review and editing, N.R., G.V.; Visualization, N.R., G.V. Supervision G.V. Project administration, P.T., G.V.; Funding acquisition, P.T. and G.V. All authors have read and agreed to the published version of the manuscript.

## Acknowledgments

G.V acknowledges support from Schmid College of Science and Technology at Chapman University for providing computing resources at the Keck Center for Science and Engineering.

