## Supplemental Figures S1-S6 for "Interpretable Atomistic Prediction and Functional Analysis of Conformational Ensembles and Allosteric States in Protein Kinases Using AlphaFold2 Adaptation with Randomized Sequence Scanning and Local Frustration Profiling"

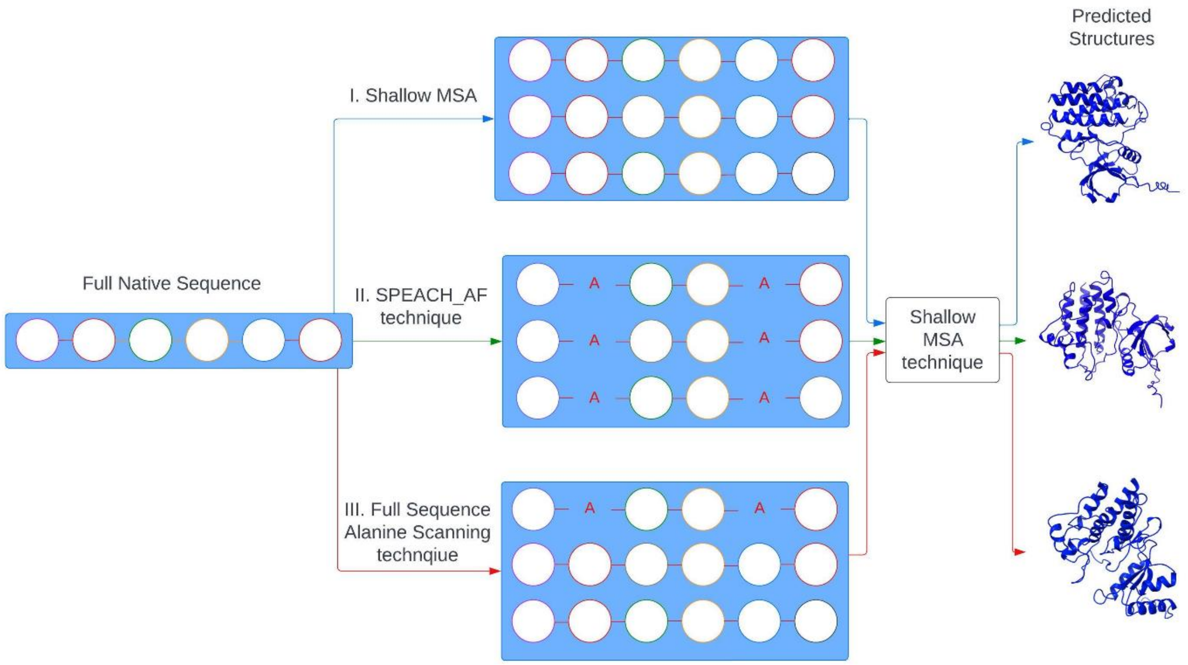


**Figure S1.** A schematic overview of the three major AF2-based adaptations used in our study : AF2 default settings, AF2 methodology with shallow MSA depth[13] ; SPEACH_AF in which the MSAs are manipulated via *in silico* mutagenesis by replacing specific residues within the MSAs[14] ; and random alanine scanning that iterates through each amino acid in the native sequence and randomly substitutes the residues to simulate random alanine substitution mutations.

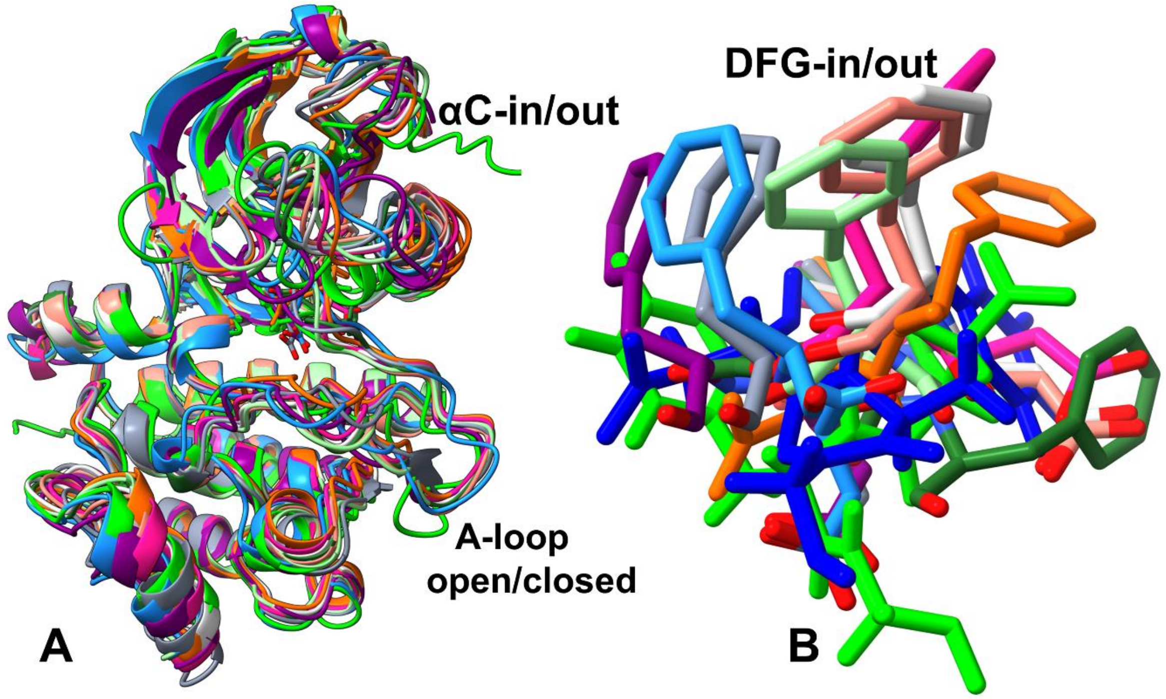


**Figure S2.** Structural alignment of the AF2-predicted conformations that are close to the inactive $I_1$ state (RMSD < 1.5 Å). (A) The predicted conformations are obtained using randomized alanine sequence scanning approach in AF2. The kinase core and the A-loop conformation shared by these models are relatively similar and closely resemble the $I_1$ state. (B) Structural alignment of the DFG motifs that adopt a number of intermediate DFG-out positions in the predicted inactive conformations. The predicted inactive states correspond to the "BLBplus " class (DFG-in/out, αC-helix-out).

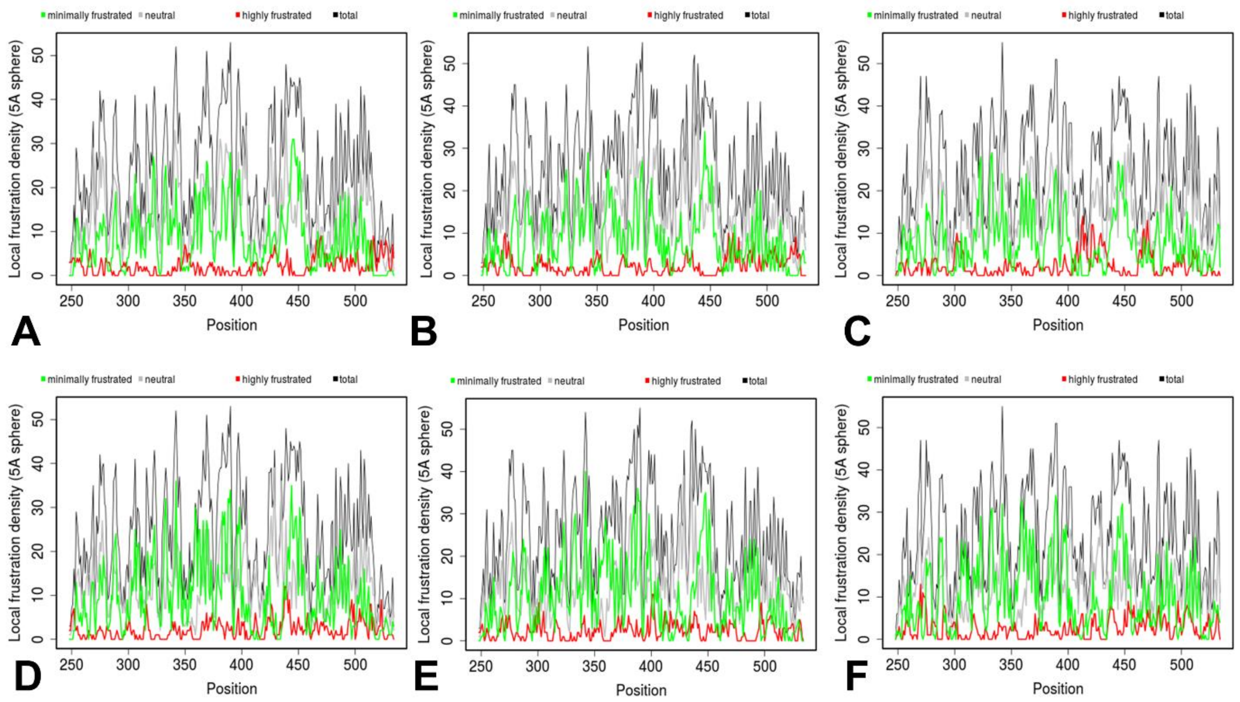


**Figure S3.** Conformational and mutational frustration analysis for the ABL states, The distributions of conformational frustration and local densities for the highly frustrated, neutrally frustrated and minimally frustrated contacts in the active ABL state (A). intermediate inactive $I_1$ state (B), and the fully inactive $I_2$ state (C). The distributions of mutational frustration and local densities for the highly frustrated, neutrally frustrated and minimally frustrated contacts in the active ABL state (D). intermediate inactive $I_1$ state (E), and the fully inactive $I_2$ state (F).

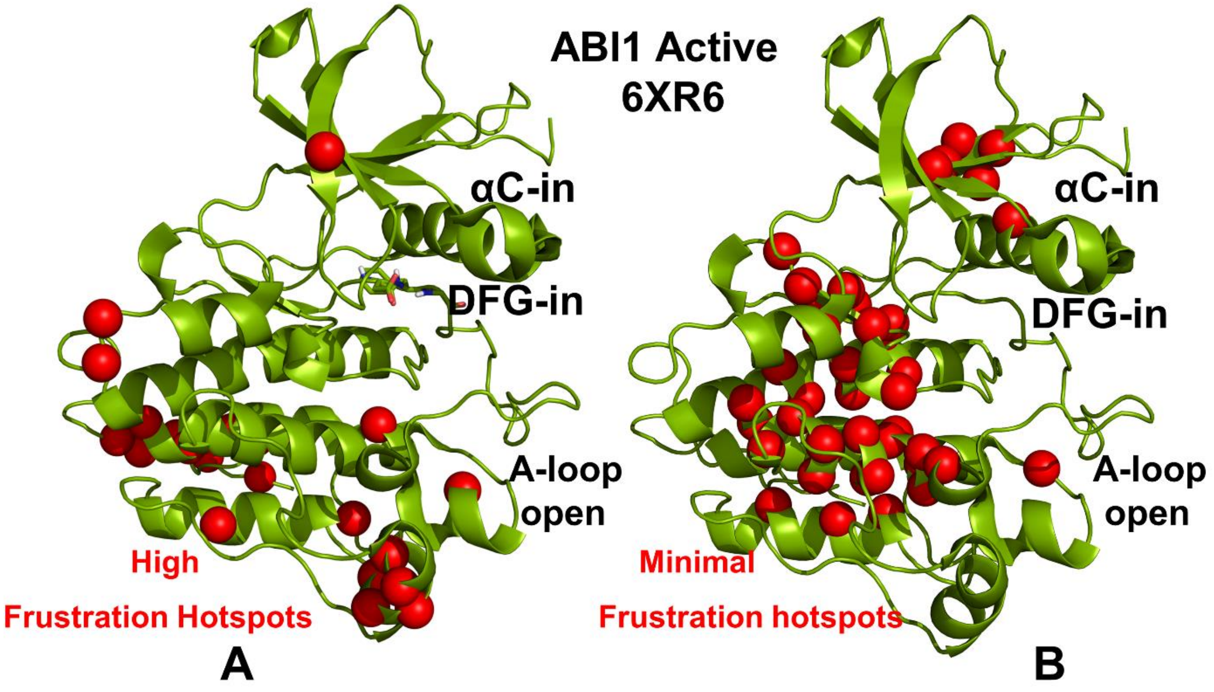


**Figure S4.** Structure-based analysis and mapping of local frustration patterns in ABL kinase. (A) Structural mapping of high frustration hotspots (the top 10% of highly frustrated sites) in the active ABL state. The kinase domain is shown in green ribbons and the high frustration hotspots are shown in red spheres. The position and conformation of the αC-helix, DFG motif and A-loop are annotated. (B) Structural mapping of minimally frustration hotspots (the top 10% of minimally frustrated sites) in the active ABL state. The kinase domain is shown in green ribbons and the minimal frustration hotspots are shown in red spheres. The position and conformation of the αC-helix, DFG motif and A-loop are annotated.

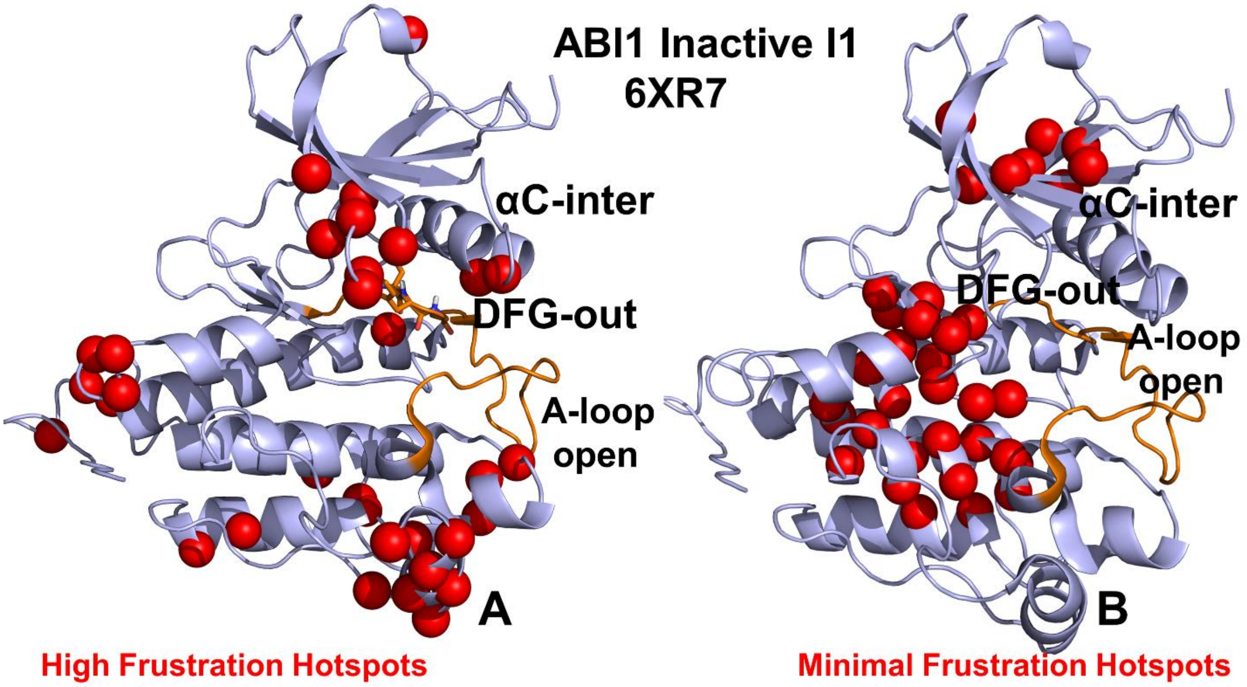


**Figure S5.** Structure-based analysis and mapping of local frustration patterns in the ABL kinase. (A) Structural mapping of high frustration hotspots (the top 10% of highly frustrated sites) in the intermediate inactive $I_1$ ABL state. The kinase domain is shown in magenta-colored ribbons and the high frustration hotspots are shown in red spheres. The position and conformation of the αC-helix, DFG motif and A-loop are annotated. (B) Structural mapping of minimally frustration hotspots (the top 10% of minimally frustrated sites) in the intermediate inactive $I_1$ ABL state. The kinase domain is shown in magenta ribbons and the minimal frustration hotspots are shown in red spheres. The position and conformation of the αC-helix, DFG motif and A-loop are annotated.

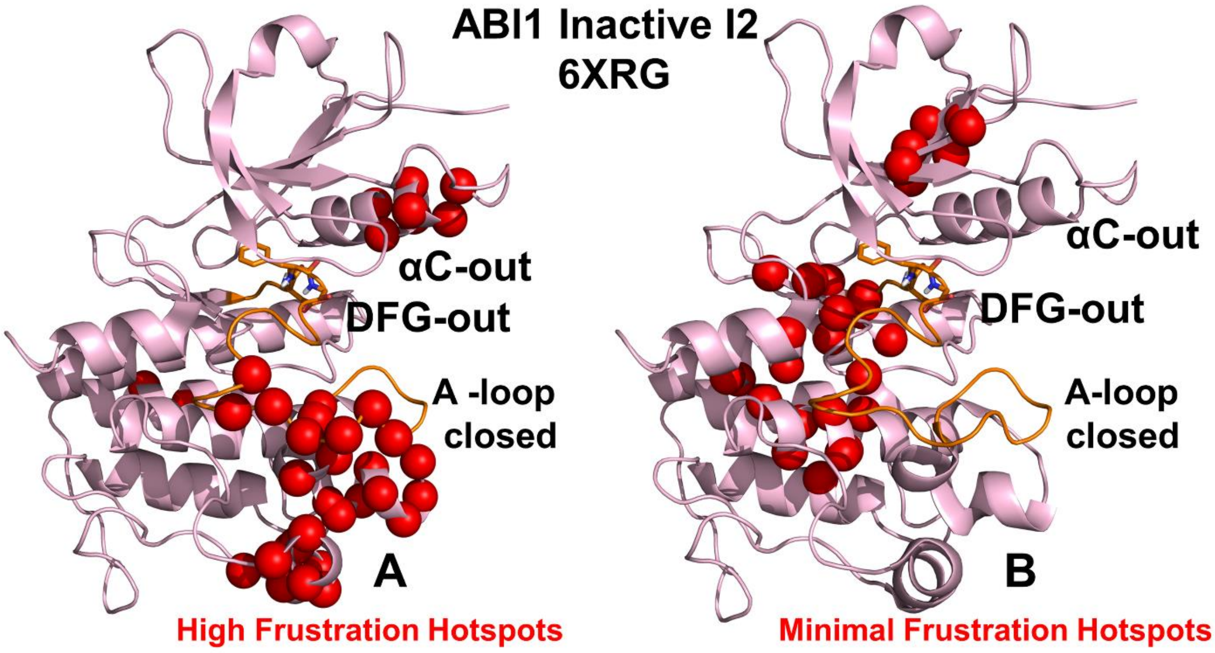


**Figure S6.** Structure-based analysis and mapping of local frustration patterns in the ABL kinase. (A) Structural mapping of high frustration hotspots (the top 10% of highly frustrated sites) in the fully inactive $I_2$ ABL state. The kinase domain is shown in light, pink-colored ribbons and the high frustration hotspots are shown in red spheres. The position and conformation of the αC-helix, DFG motif and A-loop are annotated. (B) Structural mapping of minimally frustration hotspots (the top 10% of minimally frustrated sites) in the fully inactive $I_2$ ABL state. The kinase domain is shown in light, pink-colored ribbons and the minimal frustration hotspots are shown in red spheres. The position and conformation of the αC-helix, DFG motif and A-loop are annotated.
